# Evaluation of gene knock-outs by CRISPR as potential targets for the genetic engineering of the mosquito *Culex quinquefasciatus*

**DOI:** 10.1101/2020.10.21.349704

**Authors:** Xuechun Feng, Lukas Kambic, Jared H.K. Nishimoto, Floyd A. Reed, Jai A. Denton, Jolene T. Sutton, Valentino M. Gantz

## Abstract

*Culex quinquefasciatus* mosquitoes are a globally widespread vector of several human and animal pathogens. Their biology and behavior allow them to thrive in proximity to urban areas, rendering them a constant public health threat. Their mixed bird/mammal feeding behavior further offers a vehicle for zoonotic pathogens transmission to people, and separately, poses a threat to the conservation of insular birds. The advent of CRISPR has led to the development of novel technologies for the genetic engineering of wild mosquito populations, yet research in *Culex quinquefasciatus* has been lagging compared to other disease vectors. Here we use this tool to disrupt a set of five pigmentation genes in *Culex quinquefasciatus* that, when altered, lead to visible, homozygous-viable phenotypes. We further validate this approach in separate laboratories and in two distinct strains of *Culex quinquefasciatus* that are relevant to potential future public health and bird conservation applications. We generate a double-mutant line, demonstrating the possibility of sequentially combining multiple such mutations in a single individual. Lastly, we target two loci, *doublesex* in the sex-determination pathway and *proboscipedia* a hox gene, demonstrating the flexibility of these methods applied to novel targets. Our work provides a platform of seven validated loci that could be used for targeted mutagenesis in *Culex quinquefasciatus* and the future development of genetic suppression strategies for this species. Furthermore, the mutant lines generated here could have widespread utility to the research community using this model organism, as they could be used as targets for transgene delivery, where a copy of the disrupted gene could be included as an easily-scored transgenesis marker.

## INTRODUCTION

The *Culex (Cx*.*) pipiens* species complex consists of several species and their hybrid forms, including *Cx. pipiens pipiens, Cx. pipiens pallens, Cx. pipiens molestus, Cx. quinquefasciatus, Cx. australicus*, and *Cx. globocoxitus*. While some of these species have limited distribution, *Cx. pipiens pipiens, Cx. quinquefasciatus*, and their hybrids are widespread globally and represent primary urban and suburban disease vectors. The ability of these species to produce fertile hybrids has been a source of an ongoing discussion of the taxonomy of these species [1] and has been proposed as a mechanism explaining the surprising adaptability of these mosquitoes, such as the “molestus” form of *Cx. pipiens* adapted to survive in the underground network of London [2]. Hybridization and high genetic diversity may further account for the rapid evolution of insecticide resistance across the *Culex pipiens* complex [3,4].

These mosquitoes are vectors for medically relevant RNA viruses such as West Nile (WNV), Japanese Encephalitis, Western Equine Encephalitis (WEEV), Eastern Equine Encephalitis (EEEV), Sindbis virus, and St. Louis Encephalitis virus, as well as nematodes that cause lymphatic filariasis [5]. Although fatalities in the United States due to these diseases are usually low in number, mosquitoes of the *Culex pipiens* complex are an established disease vector that represents a fertile ground for the establishment of new pathogens. It has been estimated that since its introduction in the United States in 1999, West Nile Virus hospitalizations alone amounted on average to ∼$56M per year [6]. Furthermore, the mixed feeding behavior of these mosquitoes on birds and mammals creates seasonal cycles of pathogen amplification in wild animals [7], which combined with their ability to vertically transmit some of these pathogens to their offspring [8], could lead to a periodic increase in the risk of pathogen transmission to people and the potential of unpredictable outbreaks. The zoonotic nature of these and other viruses allows for unchecked reservoirs, mostly in wild birds, that could act as evolutionary melting pots for the development of new viral forms. For example, Western Equine Encephalitis is considered to be the outcome of recombination between EEEV and a Sindbis-like virus [9]. Within this species complex, *Cx. quinquefasciatus* is considered the disease vector with the greatest human health impact worldwide, mostly due to its widespread distribution in urban and suburban areas, due to its capability of tolerating polluted water reservoirs for its larval development often associated with human and livestock populations [10,11]. Furthermore, its genetic adaptability and capability to interbreed with other species have made it a uniquely challenging vector to keep at bay.

In addition to affecting human health, invasive *Cx. quinquefasciatus* has been implicated in the decline and extinction of island avifauna due to its ability to vector avian pathogens. These effects are well characterized in the Hawai’i archipelago, where accidental introductions of *Cx. quinquefasciatus* in the mid-1800s contributed to extinctions of several Honeycreeper species (Drepanididae) and ongoing risks for those that remain [12–14], representing a serious threat to Hawaiian avian conservation. In Hawai’i and elsewhere, *Cx. quinquefasciatus* is one of the main vectors of *Plasmodium relictum*, which causes avian malaria, as well as being a competent vector for the virus causing avian pox [15]. These diseases continue to collapse endemic avian communities and their ranges [16,17] and pose an increasing concern to conservationists.

Existing genetic control tools have mainly been developed to target mosquito species belonging to the genera *Anopheles* and *Aedes*, and in the case of the latter have been implemented in the field to control human diseases [18–20]. Recently, the advent of CRISPR has fast-tracked the development of strategies for using genetically engineered vectors to modify or suppress wild populations of pests and invasive species [21,22], including technologies such as engineered gene drives put forward more than a decade ago [23]. Strategies have been developed for the engineering of natural populations, such as the use of the endosymbiont *Wolbachia* [24,25] or engineered gene drive systems. Such gene drive strategies, based on CRISPR, have been successfully tested in a variety of organisms, and proof-of-principle has been obtained in yeast [26], fruit flies [27,28], *Anopheles* [29–31] and *Aedes* [32] mosquitoes, as well as the mouse [33].

Compared to *Anopheles* and *Aedes* mosquitoes, however, *Culex* has been somewhat neglected regarding the development of such genetic strategies, and only a few studies have recently evaluated the activity of CRISPR in this species [34–36]. Although no extensive CRISPR technology development has yet been achieved, previous work has shown the feasibility of using CRISPR by disrupting the *white* [35] or *kynurenine-hydroxylase* [34] genes causing a white-eye phenotype, as well as the disruption of a CYP450 gene involved in pesticide metabolism [36]. These studies highlight the potential for the development of genetic control strategies for *Culex* mosquitoes, as well as a need to identify and characterize additional CRISPR targets and develop reliable protocols.

The scope of this study is to expand the available set of target genes that could be used in future work aimed at the development of genetic technologies for population control of *Cx. quinquefasciatus*. We chose to target a set of pigmentation and developmental genes in *Cx. quinquefasciatus* with phenotypes that have been previously evaluated in diverse species [29,30,32,37,38]. The pigmentation genes result in homozygous-viable mutations that could be used to track transgenes without the use of fluorescent markers. At the same time, the validated developmental genes could be included in future genetic suppression strategies. In this study, we include two previously validated genes (*white* and *kynurenine-hydroxylase*), three additional pigmentation genes (*cardinal, yellow*, and *ebony*), a sex-determination gene (*doublesex*), and a developmental hox gene (*proboscipedia*). We further evaluate our approach in two separate *Cx. quinquefasciatus* lines, demonstrating its flexibility of adaptation to different laboratory strains as well as the broad applications targeted to diverse genetic populations.

## RESULTS

### Targeting of *Culex quinquefasciatus* pigmentation genes for disruption

In order to validate multiple gene targets that could be used in future research, we identified homologs of the fruit fly genes *white (w), cinnabar* (also known as *kynurenine hydroxylase - kh*, or *kynurenine mono-oxygenase - kmo*), *cardinal (cd), yellow (y)*, and *ebony (e)* from *Culex quinquefasciatus*. For each gene, we first designed primers to PCR-amplify genomic sequences covering exons from our *Cx. quinquefasciatus* laboratory line sourced from California (CA line). Using the sequences obtained from such PCRs and the available genome for this species [39], we designed multiple gRNAs targeting each genes’ coding sequences (**Fig. 1**). Remarkably by sequencing the *white* gene in our line, we identified two different alleles with a quite extensive divergence, distinguishable by several single nucleotide polymorphisms (SNP) every 10-20 bp (**Supplementary information**), and decided to design one of the gRNAs (*w4*) on a position covering one of these SNPs, to evaluate how this could affect targeting. This design was chosen to evaluate whether the injection of a gRNA matching a specific allele would be able to efficiently target a sequence with SNPs present.

**Figure 1.**
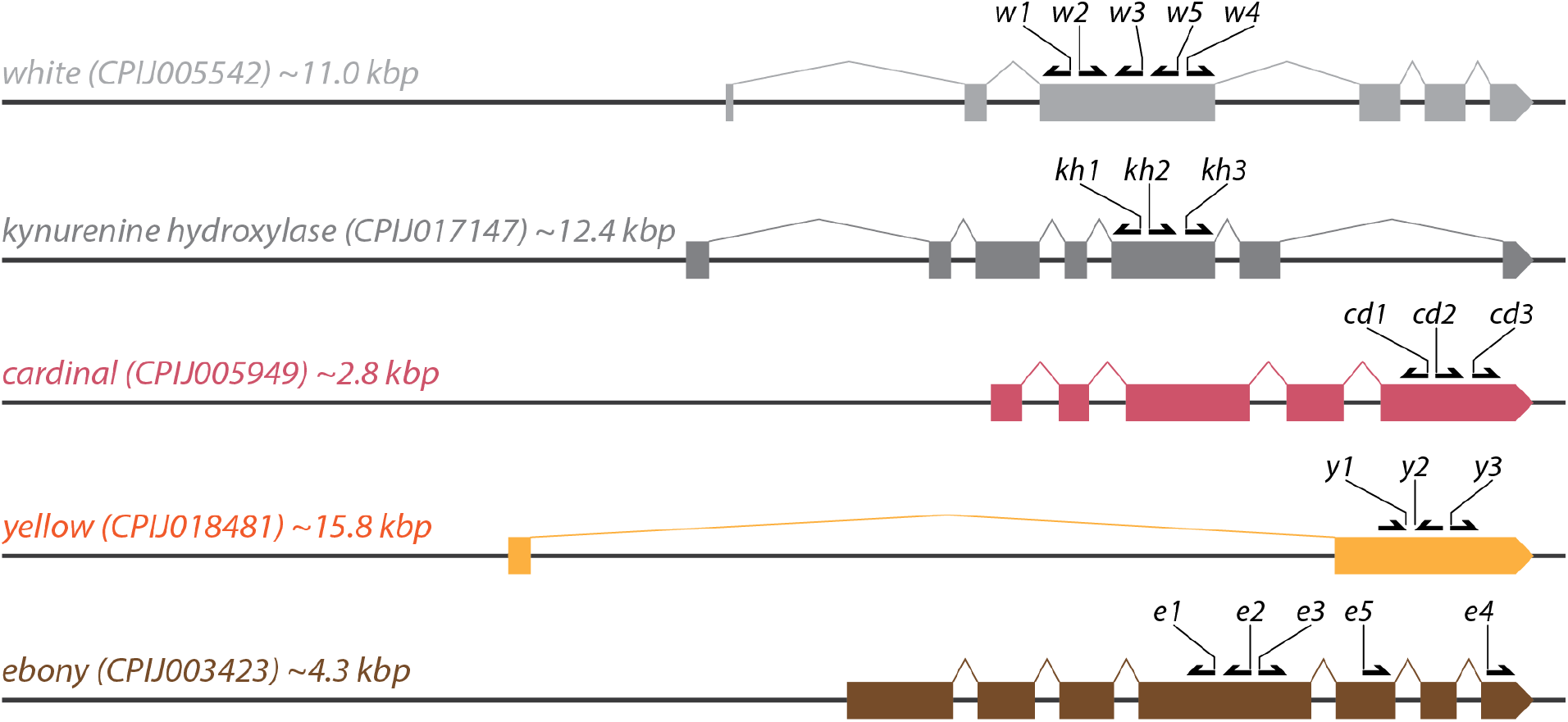
*Culex quinquefasciatus* pigmentation loci targeted. Schematic outlining the coding sequence structure of the targeted genes and the locations of the gRNA target sites used in the manuscript. The approximate locus size is indicated for each gene from the start to the stop codon. The gRNAs *w1, w2*, and *w3* were previously described as active [40] and were used here as controls. Drawings are not to scale.

We performed a preliminary test by injecting eggs with Cas9/gRNA mixes for each of the gRNAs and allowing the embryos to develop to adulthood to screen for any mosaic display of the expected phenotypes: lighter eyes for *white* (**Fig. S1**) and *kh*, red-eye for *cardinal*, lighter body for *yellow*, and darker body for *ebony*. For each gene, we observed at least one gRNA yielding the expected phenotypes, and based on this preliminary analysis. We were able to select a gRNA for each gene to carry forward for further analysis of each mutant phenotype. Interestingly, the gRNA *w4* gave comparable results to the previously validated *w3* [40], even if our injected population contained a high frequency of a SNP within the targeted sequence (**Supplementary information**).

### Isolation of *Culex quinquefasciatus* pigmentation mutants

In order to recover and characterize homozygous mutant lines, we performed an additional round of injection for the selected gRNAs, as well as for the previously validated *w3*-gRNA targeting *white* as a control [40]. For each gene, we injected ∼100-400 eggs using the same protocol from our preliminary testing and scored the mosaic phenotype in the G0 injected animal at the pupal stage (**Table 1**). For the genes *w, kh, cd* we were able to clearly identify animals displaying a mosaic phenotype (**Fig. S1**), while for the *yellow* and *ebony* the mosaicism was not as obvious. For *ebony*, we were able to identify some animals displaying a darker body phenotype (**Fig. S1**), yet we decided to score them as wild-type in **Table 1** as it was difficult to make a clear distinction. This difficulty in identifying mosaic *ebony* phenotypes suggests that *ebony* is likely to be a non-cell-autonomous phenotype, similar to the fruit fly, affecting neighboring cells with a wild-type genotype and resulting in varying gradients of the phenotype depending on the level of mosaicism.

**Table 1.** Summary of injection information and phenotypic screening. The table summarizes the injection and mutant screening information. *: G1 offspring of a G0 pool displaying wild-type phenotype. **: G1 offspring of a G0 pool displaying mosaic *kh-/kh+* phenotype. ***: For these two injections, we did not score the mosaic phenotype as it was hard to discern; therefore, some of the individuals scored as wild-type might have been mosaic. §: Used a modified protocol for this injection (see Methods). ‡: The G0 animals generating this G1 offspring were outcrossed to wild type, and therefore, these animals were not scored for the *ebony-* phenotype as it would have been heterozygous.

| injection |  |  | G0 |  |  |  | G1 | G2 |
| --- | --- | --- | --- | --- | --- | --- | --- | --- |
| Target Gene | Line | gRNAs | Injected eggs | Adult survival | Wild-type phenotype | Mosaic phenotype | Mutants/Total (%) | Mutants/Total |
| *white* | CA | *w4* | 337 | 34 (10.1%) | 3 | 31 | 174/174 (100%) | - |
| *kh* | CA | *kh3* | 400 | 46 (11.5%) | 13 | 33 | 0/389 (0%)<br>* | - |
|  |  |  |  |  |  |  | 25/25 (100%)<br>** | - |
| *cardinal* | CA | *cd1* | 290 | 32 (11.0%) | 0 | 32 | 28/84 (33.3%) | - |
| *yellow* | CA | *y1* | 220 | 25 (11.4%) | 25<br>*** | n/a<br>*** | 71/159 (44.7%) | - |
| *ebony* | CA | *e4* | 288 | 90 (31.3%) | 90<br>*** | n/a<br>*** | 0/375 (0%) | - |
|  |  | *e1 + e2 + e4* | 240 | 11 (4.6%) | 5 | 6 | 43/238 (18.1%) | - |
| *white* | HI | *w3* | 258 | 41 (15.9%) | 27 | 14 | 738/805 (91.7%) | - |
| *ebony* | HI-*white* | *e5* | 533 | 34 (6.4%)<br>§ | 15 | 19 | n/a<br>‡ | *white*: 82/558<br>*ebony*: 76/558<br>*ebony-white*: 52/558 |

To recover homozygous mutants of the targeted genes, we collected the G0 injected animals for each of *white* (*w4*), *cardinal* (*cd1*), *yellow* (*y1*), and *ebony* (*e4*), pool-crossed them to each other (one pool per gene), and scored their progeny for the expected phenotypic display. For all genes, except *ebony*, we were able to recover homozygous mutants that displayed the expected phenotypes: white eyes for *white*, red eyes for *cardinal*, and lighter body for *yellow* (**Fig. 2**). Remarkably, for *white*, 100% of the G1 progeny was mutant, suggesting very high activity of the *w4* gRNA, while for *cardinal* and *yellow*, we instead recovered 33% and 45% mutant G1, respectively (**Table 1**). Because the *kh* injection resulted in the highest G0 survival and the phenotype was easily screened, we decided to generate two separate G0 crosses by pooling together the animals displaying a mosaic phenotype and separately pooling the ones having wild-type eye color. For the *kh*-mosaic pool, we recovered 100% G1 progeny displaying the expected white-eye phenotype (**Fig. 2**), while for the *kh*-wild-type pool, we did not observe any homozygous mutant animals in the G1 progeny (**Table 1**). While our data for the *kh*-mosaic pool comes from a single raft, and thus the high rate of mutant recovery could be due to an exceptional situation in which the two parents had high germline mutagenesis, our results from the *kh*-wild-type pool suggest that animals not displaying a phenotype had poor germline targeting. This underscores the usefulness of visible markers during the development of genetic tools for population engineering, as they could help the recovery of transgenic animals.

**Figure 2.**
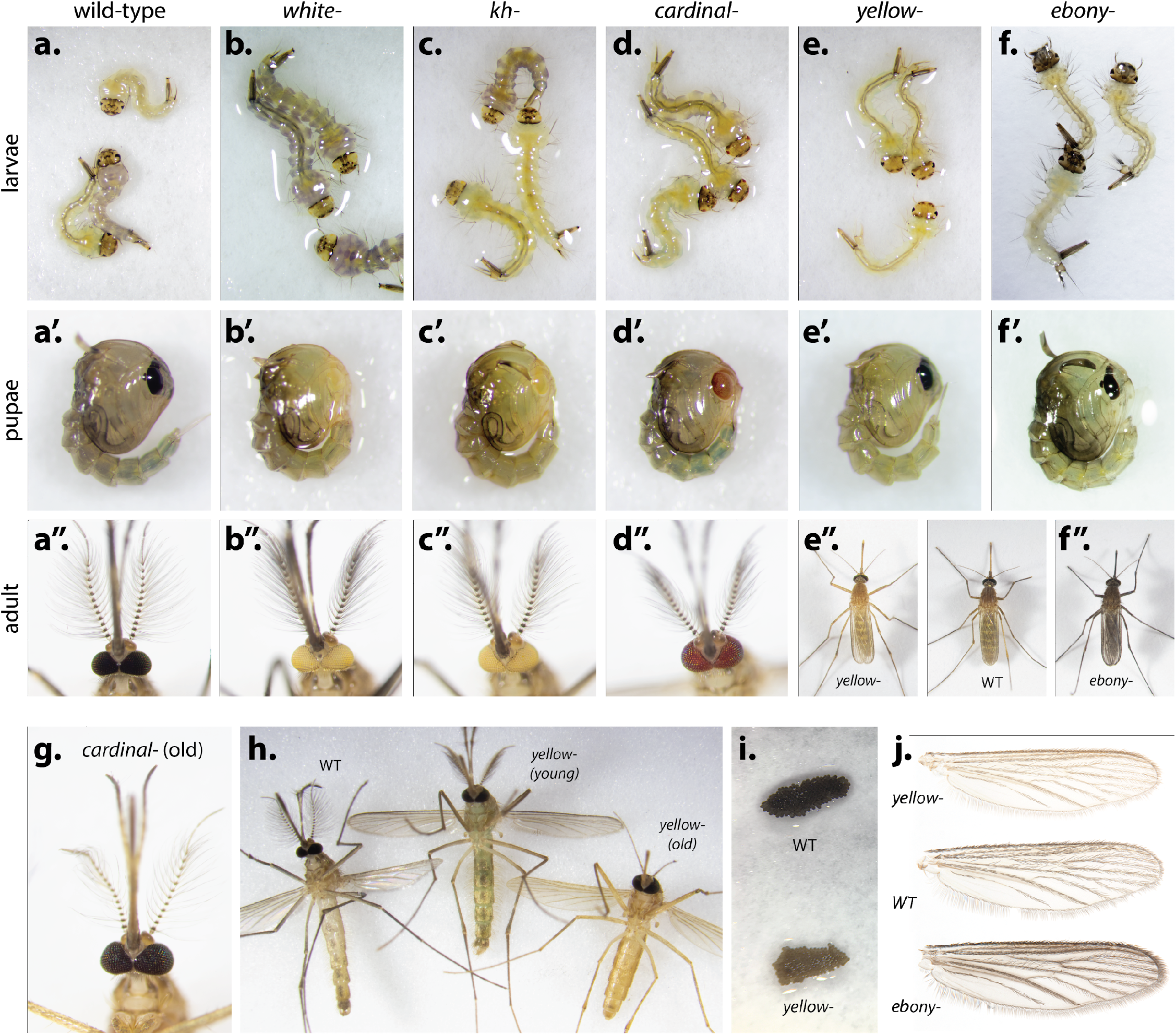
Phenotypes of the generated mutant animals generated. (**a-e**) Larvae, (**a’-e’**) pupae and (**a’’-e’’**) adults of (**a, a’, a’’**) wild-type mosquitoes and (**b, b’, b’’**) *white-*, (**c, c’, c’’**) *kynurenine-hydroxylase- (kh-)*, (**d, d’, d’’**) *cardinal-*, (**e, e’, e’’**) *yellow-*, and (**f, f’, f’’**) *ebony-* mutants. (**g**) cardinal mutants darken over time, acquiring a phenotype similar to wild-type (**a’’**). (**h**) *yellow-* mutants eclose from the pupae (center) with a phenotype similar to wild-type (left) and lighten over time to assume a more recognizable lighter color (right). (**i**) *yellow-* mutant females lay eggs that are lighter in color (left) compared to wild-type ones (right). (j) Lighter and darker wing pigmentation of *yellow-* and *ebony-* mutants, respectively, compared to a wild-type (WT) control.

As we were not able to recover G1 *ebony*- mosquitoes in our *e4* injection, we attempted to increase mutagenesis efficiency by performing an additional injection for *ebony* employing a mix of three gRNAs (*e1, e2, e4*). We noticed that the survival rate of this injection was lower than the one performed before (∼5% G0 surviving compared to ∼31% for the *e4* injection), suggesting that there might be some lethality associated with this gene or with one of the gRNAs used. Additionally, we scored 6/11 of the G0 animals as putative mosaics given their starkly darker phenotype. We then performed a pooled cross of these 11 surviving mosquitoes and obtained 43/238 G1 animals (∼18%, **Table 1**) displaying a dark *ebony-* phenotype (**Fig. 2**).

To evaluate whether our analysis would be generalizable to different *Cx. quinquefasciatus* strains, which would need to be modified in the laboratory to pursue insect control applications targeting different mosquito populations, we targeted a separate line sourced from Hawai’i (O‘ahu and Hawai‘i Islands). Here, we used the validated *w3-*gRNA, which was chosen for its high activity and obvious phenotype, as noted in [40] and this study, and performed a similar analysis to what was described above. For this line, ∼34% of G0 injected animals displayed a *white-/white+* mosaic phenotype, and an intercross of such G0 animals led to a recovery of mutants in the G1 progeny with ∼92% rate (**Table 1**), comparable to what was previously described above using *w4*, and with the same gRNA used in previous work [40]. We used the recovered G1 animals to establish a homozygous *white* mutant line (HI-*white*) to be used in the next steps of this work.

### Generation of a *white-*/*ebony*- double mutant line of *Culex quinquefasciatus*

We then decided to test whether we could reliably obtain sequential mutagenesis of different genes. We chose the HI-*white* line to evaluate whether this single-mutant colony could be further utilized for downstream studies that might require targeting the genome for transgenes. For example, in the model organism *Drosophila melanogaster*, single mutant lines are routinely used for subsequent transgenesis with synthetic constructs carrying a copy of the disrupted gene, which functions as a selectable marker for tracking integration efficiency [41]. We then subjected the HI-*white* line to an additional round of mutagenesis with the injections of the *e5*-gRNA targeting *ebony*. To avoid a potential compounded weakness of the double-mutant G0 animals, usually weaker due to the injection process, we modified our crossing strategy by backcrossing the *e5*-injected G0 mosaic individuals to wild-type. We then inter-crossed the G1 animals (all of which displayed wild-type phenotypes) to screen the G2 progeny for the *white, ebony*, and *ebony-white* phenotypes (**Fig. S2**). Indeed, we were able to recover 76/558 *ebony-* homozygous mosquitoes using this strategy, as well as obtaining 52/558 homozygous double-mutant animals that could be used as lines for future transgenesis. Based on the genetic cross utilized, these numbers suggest that *white* and *ebony* are segregating independently. Additionally, by using the phenotypes observed in the G2 offspring, we were able to estimate the amount of edited G1 that we would have observed for this experiment if the G0s were instead intercrossed (∼92%, see the Methods section for detailed calculation). This value allows us to compare the ebony-results to the other conditions and it suggests that the editing rate of the *e5*-gRNA is comparable to the levels observed for other loci.

### Phenotypical and molecular characterization of the mutants generated

As mentioned previously we were able to recover homozygous mutant mosquitoes for each of the targeted genes and establish homozygous lines. The observed phenotypes illustrated in **Fig. 2** paralleled previous work performed in the fruit fly, *Anopheles [29,37]*, and *Aedes [32]* species. We observed distinct white-eye phenotypes for *white* (**Fig. 2b**) and *kh* (**Fig. 2c**) mutants and a red-ish eye for *cardinal* mutants (**Fig. 2d**), which were visible in all stages of development but more obvious in the 4th larval instar, the pupae, and the adult. Interestingly, *cardinal-* mutants would darken after eclosing from the pupae and, over time (2-3 days), reach a phenotype indistinguishable from wild type (**Fig. 2g**) similarly to what was previously observed in *Anopheles* [37]. For mutants of the yellow gene, we could not see a major difference at the larval/pupal stages, but at the adult stages, the lighter body color phenotype becomes more obvious (**Fig. 2e-e’** vs. **2e’’**). Contrary to the *cardinal* mutants, *yellow* mosquitoes lighten over time, rendering the phenotype more obvious in older animals (**Fig. 2h**). Lastly, eggs laid by *yellow-* mosquitoes also appear lighter in color (**Fig. 2i**), probably due to the melanin pathway [42] being involved in the tanning of the eggs. For ebony, we observed an obvious phenotype in the larvae, which display distinctly darker hair, head cap, and siphon (**Fig. 2f**). The *ebony-* pupae also display a darker phenotype, although it might be difficult to distinguish ebony mutants from late-stage wild-type pupae as they darken before eclosion (**Fig. 2f’**). Adult *ebony-* animals display a dark body phenotype which can be observed on all body parts (**Fig. 2f’’**), and it is especially evident on the legs and the wings (**Fig. 2j**) when observed under the microscope.

To confirm the editing at the correct location, we isolated genomic DNA from a few mosquitoes from each of the mutant lines generated and performed PCR amplification of the targeted region. Sanger sequencing revealed the makeup of the mutations generated during the editing process. As we generated mutant lines by pooling G1 (or G2) mutant animals, we observed multiple alleles during this analysis, including mutant mosquitoes heterozygous for different alleles. In such situations, the ICE analysis tool allowed us to identify the different alleles present in the mixed Sanger traces for each of the *white* (**Fig. S3**), *kh* (**Fig. S4**), *cardinal* (**Fig. S5**), *ebony* (**Fig. S7**) lines. We observed only one allele for *yellow* (**Fig. S6**), most likely due to the establishment of the yellow line by a single raft carrying only one type of mutation. For *ebony*, we observed efficient targeting at the *e1* and *e2* locations (**Fig. S7**), while no cleavage was observed at the *e4* site. This suggests that *e4* is less efficient than the other two gRNAs and it would explain our inability to recover *ebony-* mutants in our initial injection using only the *e4*-gRNA (**Table 1**). Indeed our molecular characterization revealed that each of the chosen genes was successfully targeted for disruption. With the exception of *yellow*, all observed indels disrupt the open reading frame. Interestingly, the indel isolated for *yellow* lacks 3 bp, and therefore maintains the frame, leaving the possibility that the recovered mutant is a hypomorphic allele that maintains a partial function of *yellow*.

### Characterization of other potential targets for gene drive targeting

In order to validate our methods on genes that could be useful in future genetic control strategies, we selected two candidates, *doublesex* (*dsx*) and *proboscipedia* (*pb*). The *dsx* gene was chosen as it has been successfully used in a gene drive suppression strategy, in which disruption of the *dsxF* female isoform of the gene transcript resulted in the conversion of females to intersex sterile individuals without altering male fitness [30]. *pb* was chosen with the goal of generating altered mouthparts, as happens in the fruit fly for homozygous mutants of this gene, that would impair the female mosquitoes’ blood-feeding process.

For *dsx* we designed a gRNA that would specifically disrupt the *dsxF* isoform of the gene (**Fig. 3a**) by disrupting the splice acceptor site leading to the expression of this isoform (**Fig. 3b**). Successful editing of this sequence could either result in a null *dsx-* allele or alternatively specifically disrupt the female transcript and promote the generation of the male transcript. Either event should lead to visible morphological alteration of the female body structures. We injected 330 eggs and observed an unusually low survival rate of 2.7%. Out of the nine surviving animals, four were males, displaying a wild-type phenotype, and five females, three of which were unable to fully eclose from the pupal case (**Table 2**). One of the two surviving females displayed signs of masculinization in the maxillary palps (MP) length, which were longer than a wild-type female but not as elongated as the male ones (**Fig. 3d-f**). To confirm that these phenotypes resulted from the *dsxF* transcript’s disruption, we sequenced PCR amplicons of the region targeted by the *dsx1*-gRNA. Indeed, in every single surviving mosquito we observed efficient disruption of the *dsxF* isoform at the target site as shown by the Sanger traces which display overlapping signals starting at the cut site (**Fig. S8**), confirming efficient targeting of this gene, and its conserved role in determining sexual dimorphism.

**Table 2.** Summary of injections and phenotypic screening for doublesex and proboscipedia. This table summarizes the injection details for targeting *doublesex* and *proboscipedia*. For *dsx*, all G0 females displayed a mutant phenotype while the males appeared wild-type. * = 2 females successfully eclosed from the pupal case, while 3 died during eclosion. We considered all 5 of these females in our survival to adulthood calculation. ** = 10 females successfully eclosed from the pupal case, while 7 died during eclosion. We considered all 17 of these females in our survival to adulthood calculation. For *proboscipedia*, we also scored the G1 offspring of a G0 mass-cross.

| injection |  |  | G0 |  |  |  |  |  | G1 |  |  |  |
| --- | --- | --- | --- | --- | --- | --- | --- | --- | --- | --- | --- | --- |
| Target Gene | Line | gRNAs | Injected eggs | Adult survival | Wild-type |  | Mutant |  | Wild-type |  | Mutant |  |
|  |  |  |  |  | Females | Males | Females | Males | Females | Males | Females | Males |
| *dsx* | CA | *dsx2* | 330 | 9 (2.7%) | 0 | 4 | 2 (+3)* | 0 | - | - | - | - |
| *dsx* | HI | *dsx2* | 206 | 31 (15.0%) | 5 | 9 | 10 (+7)** | 0 | - | - | - | - |
| *pb* | CA | *pb1 + pb2 + pb3* | 250 | 13 (5.2%) | 6 | 2 | 3 | 2 | 44 | 45 | 0 | 0 |

**Figure 3.**
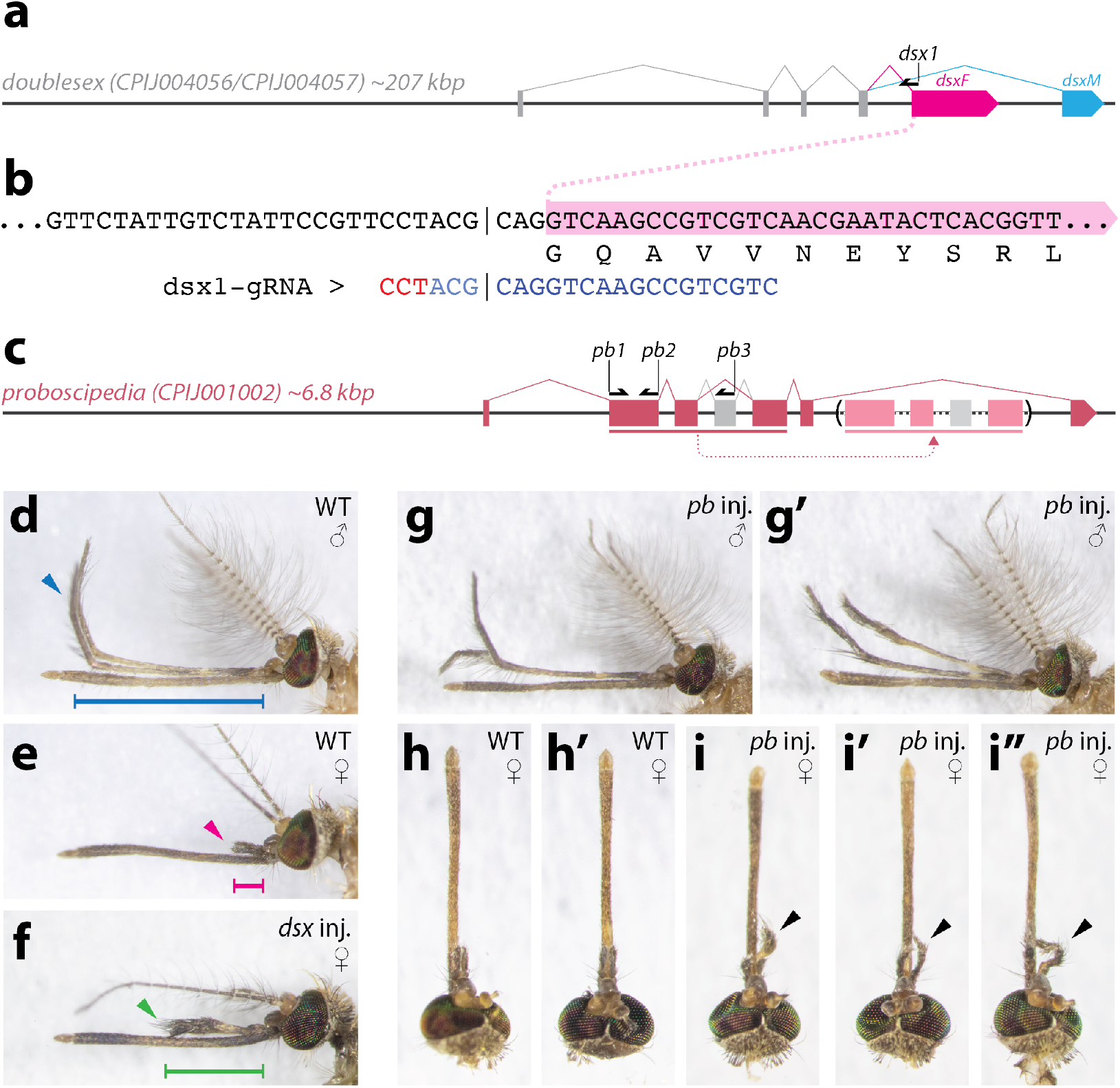
Editing of the *doublesex* and *proboscipedia* genes. **(a)** schematic of the *Culex quinquefasciatus* doublesex gene, displaying the two different splicing isoforms *dsxF* (pink) and *dsxM* (blue) characteristic of males and females. The dsx1 gRNA target was chosen on a sequence close to the *dsxF* splicing acceptor, in order to specifically disrupt this isoform. **(b)** sequence location of the *dsx1* gRNA relative to the spicing site. *dsxF* exon indicated in pink. **(c)** Structure of the *proboscipedia* gene. The annotated gene CPIJ001002 does not correspond to the genomic sequence that we have observed in our strain. Based on homology with other species, we have identified an unannotated putative exon indicated in gray. Also, the current annotation displays an exact duplication of a 1285 bp sequence indicated with horizontal bars and arrows under the gene. **(d)** Wild-type male with long maxillary palps (MP) curved upwards (blue bar and arrow). **(e)** wild-type female with short MPs (magenta bar and arrow) **(f)** *dsx*-injected G0 eclosed female showing elongated MPs (green bar and arrow). **(g-g’)** *proboscipedia*-injected males, showing altered MPs, in **(g)** one of the MPs is bent downwards, and in **(g’)** one is shorter than the other, and they appear malformed. **(h-h’)** Wild-type female heads. **(i-i’-i’’)** *pb*-injected females displaying altered MPs, slightly longer than wild-type ones and displaying leg-like joints **(h-i)** antennae removed.

To confirm the reliability of the chosen gRNA targeting *dsxF*, we also evaluated its ability to edit the gene in our Hawaii *Cx. quinquefasciatus* line. We performed an injection with a lower concentration of Cas9 in an attempt to increase survival compared to our CA line results (see Methods for details). From the 206 injected eggs, 31 (15.0%) survived to adulthood (**Table 2**). There were 9 males and 5 females displaying wild-type MPs and 17 females with elongated MPs. Seven of the 17 modified females failed to properly eclose and died shortly after. Similarly to what was observed for the CA line, the elongated MPs in G0 females resembled the males’ ones with an even more pronounced phenotype compared to what we observed in the CA G0s (**Fig. S9**). These data further support the generalization of these techniques in diverse strains and reinforce the potential use of the tested gRNA in future population suppression strategies. As *dsx-* females were expected to be sterile, we generally limited our phenotypic and molecular analysis to the G0 animals. However, using the HI line, we crossed our surviving 9 G0 males to 21 wildtype females and crossed our surviving 10 G0 *dsx-* phenotype females to 21 wildtype males. The G0 males produced a single egg raft, from which males and females hatched and developed normally. Instead, the G0 females did not produce any egg rafts after 70 days and multiple blood meal provisions, suggesting that *dsx* disruption led to G0 female sterility in this experiment.

To evaluate the potential targeting of *proboscipedia* (*pb*) we first identified the genomic location of this gene in our laboratory line. Surprisingly, our genomic sequence differed substantially from the reference genome for the *pb* locus (CPIJ001002). In comparison to our California line, the annotated *pb* gene contains duplications of exons 2, 3, and 4, which appear downstream of exon 5 (**Fig. 3c**, duplication of exons in parenthesis). Additionally, by homology with the fruit fly, we identified an additional putative exon between exons 3 and 4 (**Fig. 3c**, gray exons). Using the sequence of our laboratory strain, we designed three gRNAs to target conserved locations of this gene to ensure disruption of its function. To ensure efficient editing, we performed an injection combining the three designed gRNAs. Of the 250 injected eggs, 13 mosquitoes survived to adulthood (**Table 2**). Five such surviving G0 animals, two males, and three females displayed a mutant phenotype on the maxillary palps (MP); the males displayed malformed or downward-bent MPs (**Fig. 3g-g’**, compared to wild-type in **Fig. 3d**), while females displayed slightly elongated MPs that seemed to show leg-like articulation (**Fig. 3i-i’’**) when compared to the wild-type counterparts (**Fig. 3h-h’**). The observed phenotypes suggest that we were able to disrupt the *pb* gene and obtain phenotypes congruent with mutations described for the fruit fly *pb* gene. While we attempted the molecular characterization of the animals displaying the observed *pb-* phenotype, we were unable to obtain PCR amplification from these samples. It is possible that the duplication annotated on the reference genome is present in our laboratory line and that the simultaneous targeting with three gRNAs of such repeated sequence results in local rearrangements in the mosaic G0 animals, which renders the DNA amplification process inefficient. Our future plans include repeating these injections and attempting the recovery of homozygous *pb-* mutants from downstream generations that could confirm the effect of homozygous disruption of this gene.

## DISCUSSION

Here we build on previous work demonstrating the feasibility of a CRISPR approach for targeted mutagenesis in *Culex quinquefasciatus* [34–36], by providing an in-depth mutagenesis analysis of several pigmentation loci in this species. The analysis described herein provides information for a set of five viable loci, *white, kynurenine-hydroxylase, cardinal, yellow*, and *ebony*, that can be efficiently targeted in future genetic strategies for population suppression of these mosquitoes. As Anderson et al., we also target the *kynurenine-hydroxylase* gene for disruption [34]. While in this study we observed a higher mutant G1 rate (100%) than in the previous study (2.6%) [34], we acknowledge that the observed difference could have been positively biased due to the screening strategy that we used specifically for the *kh* gene, in which the G1 is obtained from a pool of G0s displaying mosaic phenotype. Alternatively, the discrepancy could be explained by a difference in the efficiency of the chosen gRNA. We also note that our data for such a pool was derived from a single raft that might have displayed an exceptional mutagenesis rate. For the *white* gene, we explored both a newly designed *w4-*gRNA and the *w3-*gRNA that was previously validated by Li et al. [40], and we observed higher rates of mutagenesis than previously described by them (92%-100% in this study, compared to 61%-86% in Li et al. [40]). Importantly, in several of our experiments, we were able to recover homozygous mutants at a high rate by intercrossing G0 injected animals. This suggests that this strategy could be used to target genes without a readily distinguishable phenotype to obtain mutant G1 at a high enough rate for experimentation. Additionally, such an array of validated visible phenotypes could aid future transgenesis of *Culex quinquefasciatus*, for example, by employing gene-cassettes as markers to recover the disrupted phenotypes. Such markers also bypass issues associated with fluorescent protein expression, such as auto-fluorescence or equipment requirements, and allow flexibility when multiple transgenes need to be combined.

We purposely designed one of our gRNA (*w4*) on a location where our CA line displayed heterogeneity in the population with different haplotypes. While the *w4*-gRNA would perfectly match one allele, it would have mismatches with others (**Fig. S3**). Nonetheless, we show efficient targeting of all alleles, suggesting that mutagenesis efforts in diverse lines of *Culex quinquefasciatus*, and perhaps other organisms, could tolerate to some extent allelic differences in a genetically heterogeneous population. In addition, this finding could have implications for gene drive strategies using CRISPR, as genetic diversity in wild populations has been considered a potential hindrance to gene drive spread. Our data suggest that this issue could be more limited than previously thought [43] and could allow for widening the pool of genetic targets available [44]. Additionally, this suggests caution for strategies aiming to use fixed alleles to contain the spread of gene drives [45], potentially restricting the pool of available targets and underscoring the need for extensive experimental validation.

In addition to the pigmentation genes targeted in this work, we evaluate the phenotypes associated with two developmental genes, *doublesex* in the sex-determination pathway, and *proboscipedia*, a hox gene involved in the differentiation of mouthparts. Our analysis suggests that the methods described herein could be used to obtain a high degree of mutagenesis for other non-viable genes or ones having pleiotropic effects. Based on the injection of Cas9/gRNA complexes in developing embryos, our techniques could be readily used for scientific inquiry needing the generation of functional knock-outs. Our work targeting the female isoform of *dsx* underscores this gene’s potential for its use in *Culex quinquefasciatus* population suppression efforts, using strategies such as the CRISPR-based gene drive explored in *Anopheles* [30].

Furthermore, our analysis validates the findings in two separate lines isolated from different geographical locations. We conducted gRNA-target validation of two of the genes in two independent laboratories using diverse colony strains, thus demonstrating the robustness of the techniques described herein. One of our colonies was sourced from the continental United States (California) in the 1950s, while the second was obtained from invasive populations in Hawai’i and established in the lab in 2016, with subsequent periodic collections from the wild. The validation of multiple targets for future work in diverse strains adds further confidence that laboratory findings aimed at the control of these mosquitoes could be widely applied to locations where the genetic background might differ from the laboratory strain. While both the lines tested in this work could be targets of population engineering aimed at public health applications, our employment of a line recently sourced from Hawai’i underscores the technical feasibility of population engineering strategies aimed at bird conservation on these islands.

Our analysis furthermore shows that it is possible to target multiple genes sequentially, an aspect that could be valuable in downstream approaches requiring multiple marked transgenes. We show that it is possible to obtain and maintain a *white-, ebony-* double mutant line, suggesting that other combinations of the mutants described here could be generated. We observed that some of the mutant lines identified here seem slightly weaker than their wild-type counterparts, although there were no major hurdles in rearing them, at least for the first few generations. In this study, we did not perform further analysis of the isolated strains, such as longevity, fecundity, or fertility, as this lies beyond this manuscript’s scope, which focused on validating novel CRISPR targets valuable for future genetic engineering efforts in this species. Indeed, our work provides a set of validated genes that could be used in future *Culex quinquefasciatus* research to deliver targeted transgenesis. These findings help pave the way toward developing population engineering technology for the control of this disease vector.

## ACKNOWLEDGMENTS

We thank members of the Sutton and Gantz laboratories for comments and edits on the manuscript. We thank Anton Cornell and Nannan Liu for kindly providing *Culex quinquefasciatus* strains used in this study.

## AUTHORS CONTRIBUTIONS

All authors contributed to the project planning and the design of the experiments. X.F., L.K., and J.H.K.N. performed the experiments and contributed to the collection and analysis of data. X.F, J.T.S., and V.M.G. wrote the manuscript. All authors edited the manuscript.

## METHODS

### University of California San Diego Protocols

#### UCSD Mosquito strain

The *Culex quinquefasciatus* CA strain was kindly provided by Anton Cornell (UC Davis); the line was originally collected near the city of Merced, California in the 1950s. It was reared at 27 ± 1°C, 75% humidity, and a 12h light/dark cycle in the insectary room at the University of California, San Diego. The adults were fed with 10% sugar water. After mating, females were fed with defibrinated chicken blood (Colorado Serum Company, USA, Cat. # 31142) using the Hemotek blood-feeding system. Egg rafts were collected 4 days after blood feeding for microinjection. Larvae were fed with fish food pellets (Blue Ridge Fish Hatchery).

#### UCSD sgRNA design and synthesis

The sgRNA was designed using CHOPCHOP [46] and synthesized according to the protocol by Krisler et al. [47]. Briefly, the double-stranded DNA template for each sgRNA was synthesized by performing PCR with a specific forward primer for each sgRNA and a common reverse primer. The PCR product was later purified with QIAquick Gel Extraction Kit (Qiagen, Cat. # 28704). sgRNAs were then synthesized with the MEGAscriptTM T7 transcription kit (Thermo Fisher Scientific, Cat. # AM1334) by performing in vitro transcription reaction with a purified DNA template incubated at 37°C overnight (12-16 hours). The MEGAClear Kit (Thermo Fisher Scientific, Cat. # AM1908) was later used to purify the in vitro transcription products. The synthesized sgRNAs were diluted to 1000ng/uL, aliquoted, and stored at −80°C.

#### UCSD Culex quinquefasciatus egg microinjection

- *Cas9/sgRNA injection mix*. The recombinant Cas9 protein (DNA Bio Inc., Cat # CP01) was dissolved in nuclease-free water, aliquoted, and stored at −80°C until use. The synthesized sgRNA was mixed with Cas9 protein (with a final concentration of 200ng/uL for each) and incubated at room temperature for 15 minutes before injection.
- *Eggs preparation for microinjection*. Four days after blood feeding, the mosquito cage was put in a dark place with an oviposition cup inside. After 30 minutes, the freshly laid egg rafts were collected and put into a Petri dish covered with a wet filter paper. Single eggs were separated from rafts and aligned with a paintbrush under a dissecting microscope. The aligned eggs were then fixed on a coverslip with double-sided sticky tape. A mixture of 700 and 27 halocarbon oil (1:1) was used to cover the aligned eggs to prevent dehydration. The cover slides with aligned eggs were fixed on glass slides and put under an inverted microscope (Olympus, BX43) for microinjection.
- *Needles preparation*. Injections were performed with Aluminosilicate glass (Sutter Instrument, Cat. # AF100-64-10) pulled by a micropipette puller (Sutter Instrument, P-1000) with program: Heat=516; Pull=65; Vel.=65; Delay=47; Pressure=500; Ramp=514. Before injection, the tips of needles were slightly opened by a Beveler (Sutter Instrument, BV-10) and then loaded with injection reagents by micro-loader (Eppendorf)
- *Microinjection*. All injections were operated in a microinjection station equipped with a FemtoJet 4 microinjector (Eppendorf). The prepared Cas9/sgRNA mixtures were injected into the posterior end of eggs to ensure efficient targeting of the germline. All injections were operated in a microinjection station equipped with a FemtoJet 4 microinjector (Eppendorf).
- *Embryos and larvae rearing after injection*. After injection, the aligned eggs were rinsed with DI water and gently removed from the double-sided tapes with a paintbrush. The eggs were then transferred to a cup with clean DI water and incubated at 27C for 36 hours until they began to hatch. Transfer all the hatched larvae to a larvae-rearing tray and feed them with fish food pallets.

#### UCSD Phenotype screening and crossing

The phenotypes of G0 and G1 were screened under the dissecting microscope at the pupae stage. For *cardinal, white*, and *kh* phenotype, the G0s with mosaic phenotype were pooled, while for *yellow* and *ebony*, with their mosaic phenotypes are hard to distinguish from G0, we pooled all the injected G0 together and screened the homozygous phenotype from G1. All the homozygous individuals from G1 were pooled to establish the homozygous colony. For yellow and ebony mutants, the heterozygous siblings were sibling crossed and the homozygous individuals were later screened from G2 to build the homozygous mutant lines.

#### UCSD Genotyping

To molecularly characterize the knockout phenotypes, the genomic DNAs were extracted from ∼10 individuals of each homozygous mutant line using a single-fly DNA extractions protocol described by Gloor et al. [48]. We added 50 uL of ddH2O to diluted extracted DNA samples to a final volume of 100 uL and used 1 uL of each sample as a template to run a 25 uL PCR reaction. The primers used for PCR amplification are designed based on the genomic sequences of genes identified in VectorBase.org. Oligos information is listed in the table below.

#### UCSD Primers used for sgRNA synthesis and PCR amplification

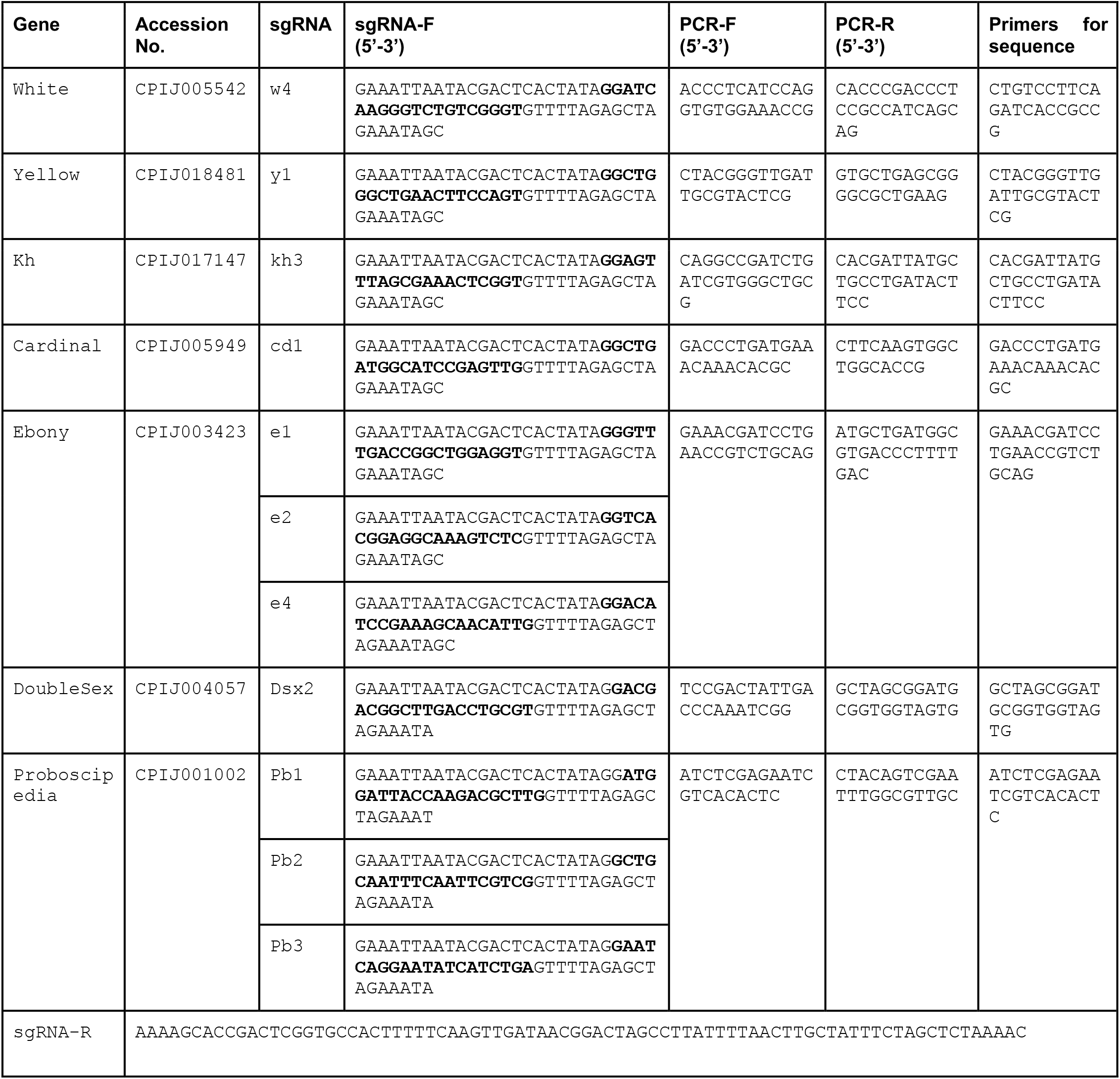

### University of Hawai’i at Hilo (UHH) Protocols

*UHH Mosquito Strain* The Hawai‘i *Culex quinquefasciatus* strain was initially collected from Hilo and Honolulu in 2016 and reared at 26 ± 1°C, 70% humidity, and a 12h light/dark cycle that includes dawn and dusk in the insectary in University of Hawai‘i at Hilo, Hilo. Adults were maintained on 3% sugar water. After mating, adults were provided commercially available bovine blood treated with sodium citrate (Lampire Biological Laboratories). Egg rafts for microinjection were collected after 1-3 opportunities for blood-feeding. Larvae were fed with fish food pellets.

#### UHH sgRNA design

A crRNA targeting *white* (w5 - GGACAGCGCGUUCAACAGGG) was based on *w*sgRNA-3 from [34] (target site on exon 3). A crRNA sequence targeting *ebony* exon 5 (e5 - CAGAGCAACUUCUACGAGCU) was selected manually and verified with CHOPCHOP and NCBI Blast. crRNA, tracrRNA, and Alt-R S. p. Cas9 Nuclease V3 were purchased from IDT.

Full *w5* crRNA as ordered from IDT: GGACAGCGCGUUCAACAGGGGUUUUAGAGCUAUGCU

Full *e5* crRNA as ordered from IDT: CAGAGCAACUUCUACGAGCUGUUUUAGAGCUAUGCU

#### UHH Culex quinquefasciatus egg microinjection

- *Cas9/sgRNA injection mix. Cas9/sgRNA injection mix*. crRNA and tracrRNA were eluted in a duplex buffer (100 mM Potassium Acetate, 30 mM HEPES, pH 7.5) to 10 pm/ul. 14.5 ul of each stock was combined, heated to 94C for 2 minutes in a thermocycler, and allowed to cool to RT. For experiments targeting *white* and *ebony*, 10 ug of Cas9 was added to each 29 ul RNA aliquot tube for a final Cas9 concentration of 330 ng/ul. This Cas9 concentration produced high developmental mortality in injections with *dsx2* (0/75 injected eggs surviving to adult). Reduction to 70 ng/uL Cas9 markedly improved survival through pupation (Table 2) and all adult UHH *doublesex* mutants were produced with this formulation. Injection mixtures were maintained at RT for 10 minutes prior to loading into Femtotip II microinjection capillaries.
- *Eggs preparation for microinjection*. Oviposition cups were placed in mosquito cages at least four days after blood feeding. After a minimum of 60 minutes, newly laid egg rafts were collected. Single eggs were separated from rafts and aligned against the edge of moistened filter paper on glass slides. No halocarbon oil was employed and all eggs were injected prior to full melanization.
- *Microinjection*. Microinjection took place under a non-inverted light microscope with a Femtojet 4i microinjector and Femtotip II microinjection capillaries (Eppendorf). The prepared Cas9/sgRNA mixtures were injected into the posterior end of the embryos to ensure efficient targeting of the germline.
- *Embryos and larvae rearing after injection*. Slides with injected eggs were gently placed into larvae trays in such a way that the slides rested against the side of the tray at ∼60-degree angles and the water’s surface just reached the place where the eggs were aligned, with the filter paper wicking water up toward the eggs and allowing them to detach from the slide surface into the water. After hatching, larvae were maintained on fish food pellets in the same insectary as the adult cages.

### Phenotype screening and crossing

Phenotypes were screened under a dissecting microscope in fourth instar larvae and/or pupae. For *white*, G0s with mosaic phenotype were pooled, and resulting homozygous G1s were pooled to establish the homozygous colony. G0 *ebony* males were separated from *ebony* G0 females and backcrossed to wildtype of the opposite sex (note that *ebony* G0s can be difficult to distinguish from wildtype). All G1s were presumed heterozygous and had fully wild-type phenotypes. G1s were pooled together to generate G2s with four different phenotypes: wildtype, *ebony, white*, and *ebony-white*.

### Calculation of the hypothetical phenotypic ratio from the G1 *ebony-* data

To calculate the editing rate for *ebony*, we are using the total of the *ebony-* animals (*ebony-*: 76/558 *ebony*-/*white*-: 52/558) which is 128/558 or ∼22.9%. To calculate the percentage of *ebony-* alleles in the G1 we generate the square root of 0.229 which is 0.479. Given that, for this experiment, we crossed the G0 to wild-type and for the other conditions we instead intercrossed the G0s, we wanted to obtain an editing value that would be comparable to the other conditions. In the G1, 0.479 is the frequency of *ebony*- edited alleles, 0.5 is the frequency of *ebony+* coming from the wild-type parent, and the remaining 0.021 are the *ebony+* alleles remaining unedited during the injection. If the G0 was intercrossed instead of crossed out to wild-type, the G0 editing rate would have been 0.479*2 = 0.958, and the expected phenotype in the G1 would have been 0.958^2 = 0.9177 ∼92%.

### Genotyping

Molecular characterization of UHH mutants was done by PCR detection of alterations in the typical Cas9 target region. Four *e5*-injected G0 individuals showed little or no amplification using a primer pair with one 3’ end 4bp upstream of the PAM site (GAGCAACTTCTACGAGC-F, TCAATGTTGCTTTCGGATGTC-R) while showing strong amplification with a control reaction, indicating homozygous editing near the target PAM site.

### ICE analysis

The analysis of mixed sequencing traces was performed by using the ICE analysis software by Synthego (Synthego Performance Analysis, ICE Analysis. 2019. v2.0. Synthego; 15 Oct. 2020), using a Sanger sequencing read performed on wild-type as a control.

## REFERENCES

1. Harbach RE. 2012 Culex pipiens: Species Versus Species Complex – Taxonomic History and Perspective. Journal of the American Mosquito Control Association. 28, 10–23. (doi:10.2987/8756-971x-28.4.10)

2. Byrne K, Nichols RA. 1999 Culex pipiens in London Underground tunnels: differentiation between surface and subterranean populations. Heredity 82 (Pt 1), 7–15.

3. Liu H, Cupp EW, Micher KM, Guo A, Liu N. 2004 Insecticide Resistance and Cross-Resistance in Alabama and Florida Strains ofCulex quinquefaciatus. Journal of Medical Entomology. 41, 408–413. (doi:10.1603/0022-2585-41.3.408)

4. Kothera L, Nelms BM, Reisen WK, Savage HM. 2013 Population genetic and admixture analyses of Culex pipiens complex (Diptera: Culicidae) populations in California, United States. Am. J. Trop. Med. Hyg. 89, 1154–1167.

5. Farajollahi A, Fonseca DM, Kramer LD, Marm Kilpatrick A. 2011 ‘Bird biting’ mosquitoes and human disease: a review of the role of Culex pipiens complex mosquitoes in epidemiology. Infect. Genet. Evol. 11, 1577–1585.

6. Staples JE, Shankar MB, Sejvar JJ, Meltzer MI, Fischer M. 2014 Initial and long-term costs of patients hospitalized with West Nile virus disease. Am. J. Trop. Med. Hyg. 90, 402–409.

7. Andreadis TG, Anderson JF, Vossbrinck CR, Main AJ. 2004 Epidemiology of West Nile Virus in Connecticut: A Five-Year Analysis of Mosquito Data 1999–2003. Vector-Borne and Zoonotic Diseases 4, 360–378.

8. Dohm DJ, Sardelis MR, Turell MJ. 2002 Experimental Vertical Transmission of West Nile Virus byCulex pipiens(Diptera: Culicidae) : Table 1. Journal of Medical Entomology. 39, 640–644. (doi:10.1603/0022-2585-39.4.640)

9. Hahn CS, Lustig S, Strauss EG, Strauss JH. 1988 Western equine encephalitis virus is a recombinant virus. Proc. Natl. Acad. Sci. U. S. A. 85, 5997–6001.

10. Fonseca DM, Smith JL, Wilkerson RC, Fleischer RC. 2006 Pathways of expansion and multiple introductions illustrated by large genetic differentiation among worldwide populations of the southern house mosquito. Am. J. Trop. Med. Hyg. 74, 284–289.

11. Okiwelu SN, Noutcha MAE. 2012 Breeding sites of Culex quinquefasciatus (Say) during the rainy season in rural lowland rainforest, Rivers State, Nigeria. Public Health Research 2, 64–68.

12. Atkinson CT, Woods KL, Dusek RJ, Sileo LS, Iko WM. 1995 Wildlife disease and conservation in Hawaii: pathogenicity of avian malaria (Plasmodium relictum) in experimentally infected iiwi (Vestiaria coccinea). Parasitology 111 Suppl, S59–69.

13. Warner RE. 1968 The Role of Introduced Diseases in the Extinction of the Endemic Hawaiian Avifauna. Condor 70, 101–120.

14. Van Riper C III, Van Riper SG, Goff ML, Laird M. 1986 The epizootiology and ecological significance of malaria in Hawaiian land birds. Ecol. Monogr. 56, 327–344.

15. Ruiz-Martínez J et al. 2016 Prevalence and Genetic Diversity of Avipoxvirus in House Sparrows in Spain. PLoS One 11, e0168690.

16. Paxton EH, Camp RJ, Gorresen PM, Crampton LH, Leonard DL Jr, VanderWerf EA. 2016 Collapsing avian community on a Hawaiian island. Sci Adv 2, e1600029.

17. Fortini LB, Vorsino AE, Amidon FA, Paxton EH, Jacobi JD. 2015 Large-Scale Range Collapse of Hawaiian Forest Birds under Climate Change and the Need 21st Century Conservation Options. PLOS ONE. 10, e0140389. (doi:10.1371/journal.pone.0140389)

18. Lacroix R et al. 2012 Open field release of genetically engineered sterile male Aedes aegypti in Malaysia. PLoS One 7, e42771.

19. Harris AF et al. 2012 Successful suppression of a field mosquito population by sustained release of engineered male mosquitoes. Nat. Biotechnol. 30, 828–830.

20. Carvalho DO, McKemey AR, Garziera L, Lacroix R, Donnelly CA, Alphey L, Malavasi A, Capurro ML. 2015 Suppression of a Field Population of Aedes aegypti in Brazil by Sustained Release of Transgenic Male Mosquitoes. PLoS Negl. Trop. Dis. 9, e0003864.

21. Gantz VM, Akbari OS. 2018 Gene editing technologies and applications for insects. Curr Opin Insect Sci 28, 66–72.

22. Esvelt KM, Smidler AL, Catteruccia F, Church GM. 2014 Emerging technology: concerning RNA-guided gene drives for the alteration of wild populations. Elife 3, e03401.

23. Burt A. 2003 Site-specific selfish genes as tools for the control and genetic engineering of natural populations. Proc. Biol. Sci. 270, 921–928.

24. Ant TH, Herd C, Louis F, Failloux AB, Sinkins SP. 2020 Wolbachia transinfections in Culex quinquefasciatus generate cytoplasmic incompatibility. Insect Molecular Biology. 29, 1–8. (doi:10.1111/imb.12604)

25. Atkinson CT, Watcher-Weatherwax W, LaPointe DA. 2016 Genetic diversity of Wolbachia endosymbionts in Culex quinquefasciatus from Hawaii, Midway Atoll, and Samoa.

26. DiCarlo JE, Chavez A, Dietz SL, Esvelt KM, Church GM. 2015 Safeguarding CRISPR-Cas9 gene drives in yeast. Nat. Biotechnol. 33, 1250–1255.

27. Gantz VM, Bier E. 2015 Genome editing. The mutagenic chain reaction: a method for converting heterozygous to homozygous mutations. Science 348, 442–444.

28. Kandul NP, Liu J, Sanchez C HM, Wu SL, Marshall JM, Akbari OS. 2019 Transforming insect population control with precision guided sterile males with demonstration in flies. Nat. Commun. 10, 84.

29. Gantz VM, Jasinskiene N, Tatarenkova O, Fazekas A, Macias VM, Bier E, James AA. 2015 Highly efficient Cas9-mediated gene drive for population modification of the malaria vector mosquito Anopheles stephensi. Proc. Natl. Acad. Sci. U. S. A. 112, E6736–43.

30. Kyrou K, Hammond AM, Galizi R, Kranjc N, Burt A, Beaghton AK, Nolan T, Crisanti A. 2018 A CRISPR-Cas9 gene drive targeting doublesex causes complete population suppression in caged Anopheles gambiae mosquitoes. Nat. Biotechnol. 36, 1062–1066.

31. Hammond A et al. 2016 A CRISPR-Cas9 gene drive system targeting female reproduction in the malaria mosquito vector Anopheles gambiae. Nat. Biotechnol. 34, 78–83.

32. Li M, Bui M, Yang T, Bowman CS, White BJ, Akbari OS. 2017 Germline Cas9 expression yields highly efficient genome engineering in a major worldwide disease vector, Aedes aegypti. Proc. Natl. Acad. Sci. U. S. A. 114, E10540–E10549.

33. Grunwald HA, Gantz VM, Poplawski G, Xu X-RS, Bier E, Cooper KL. 2019 Super-Mendelian inheritance mediated by CRISPR–Cas9 in the female mouse germline. Nature 566, 105–109.

34. Anderson ME, Mavica J, Shackleford L, Flis I, Fochler S, Basu S, Alphey L. 2019 CRISPR/Cas9 gene editing in the West Nile Virus vector, Culex quinquefasciatus Say. PLoS One 14, e0224857.

35. Li M, Li T, Liu N, Raban RR, Wang X, Akbari OS. 2019 Methods for the generation of heritable germline mutations in the disease vector Culex quinquefasciatus using clustered regularly interspaced short palindrome repeats-associated protein 9. Insect Molecular Biology. (doi:10.1111/imb.12626)

36. Itokawa K, Komagata O, Kasai S, Ogawa K, Tomita T. 2016 Testing the causality between CYP9M10 and pyrethroid resistance using the TALEN and CRISPR/Cas9 technologies. Scientific Reports. 6. (doi:10.1038/srep24652)

37. Carballar-Lejarazú R et al. 2020 Next-generation gene drive for population modification of the malaria vector mosquito, Anopheles gambiae. Proc. Natl. Acad. Sci. U. S. A. 117, 22805–22814.

38. Port F, Chen H-M, Lee T, Bullock SL. 2014 Optimized CRISPR/Cas tools for efficient germline and somatic genome engineering in Drosophila. Proc. Natl. Acad. Sci. U. S. A. 111, E2967–76.

39. Arensburger P et al. 2010 Sequencing of Culex quinquefasciatus establishes a platform for mosquito comparative genomics. Science 330, 86–88.

40. Li M, Li T, Liu N, Raban RR, Wang X, Akbari OS. 2020 Methods for the generation of heritable germline mutations in the disease vector Culex quinquefasciatus using clustered regularly interspaced short palindrome repeats-associated protein 9. Insect Molecular Biology. 29, 214–220. (doi:10.1111/imb.12626)

41. Qian S, Varjavand B, Pirrotta V. 1992 Molecular analysis of the zeste-white interaction reveals a promoter-proximal element essential for distant enhancer-promoter communication. Genetics 131, 79–90.

42. Wittkopp PJ, Stewart EE, Arnold LL, Neidert AH, Haerum BK, Thompson EM, Akhras S, Smith-Winberry G, Shefner L. 2009 Intraspecific polymorphism to interspecific divergence: genetics of pigmentation in Drosophila. Science 326, 540–544.

43. Drury DW, Dapper AL, Siniard DJ, Zentner GE, Wade MJ. 2017 CRISPR/Cas9 gene drives in genetically variable and nonrandomly mating wild populations. Science Advances. 3, e1601910. (doi:10.1126/sciadv.1601910)

44. Schmidt H, Collier TC, Hanemaaijer MJ, Houston PD, Lee Y, Lanzaro GC. 2020 Abundance of conserved CRISPR-Cas9 target sites within the highly polymorphic genomes of Anopheles and Aedes mosquitoes. Nat. Commun. 11, 1425.

45. Sudweeks J et al. 2019 Locally Fixed Alleles: A method to localize gene drive to island populations. Sci. Rep. 9, 15821.

46. Labun K, Montague TG, Gagnon JA, Thyme SB, Valen E. 2016 CHOPCHOP v2: a web tool for the next generation of CRISPR genome engineering. Nucleic Acids Res. 44, W272–6.

47. Kistler KE, Vosshall LB, Matthews BJ. 2015 Genome engineering with CRISPR-Cas9 in the mosquito Aedes aegypti. Cell Rep. 11, 51–60.

48. Gloor GB, Preston CR, Johnson-Schlitz DM, Nassif NA, Phillis RW, Benz WK, Robertson HM, Engels WR. 1993 Type I repressors of P element mobility. Genetics 135, 81–95.

