## Supplementary Information for "Evaluation of gene knock-outs by CRISPR as potential targets for the genetic engineering of the mosquito *Culex quinquefasciatus*"

**Figure S1**

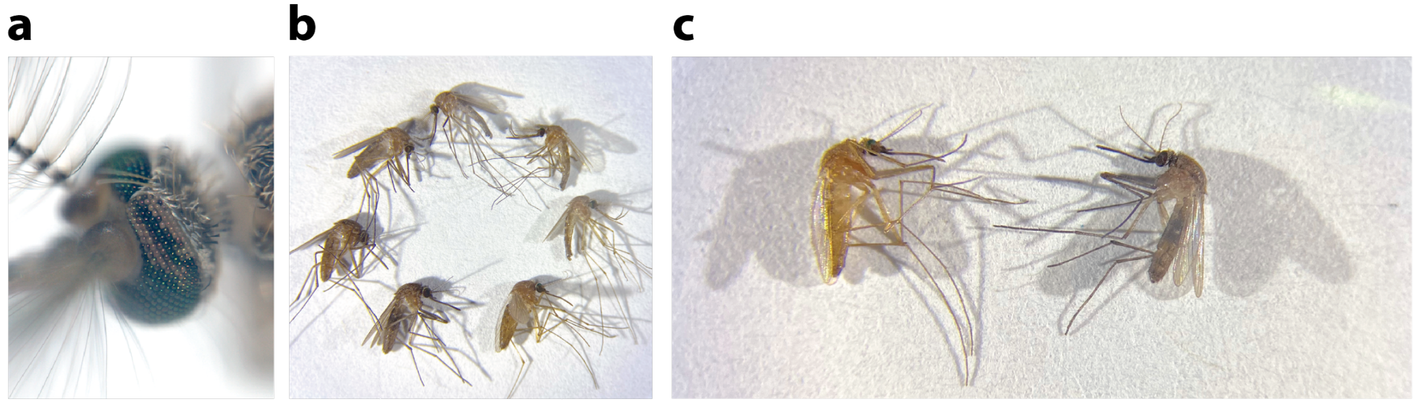


**Figure S1 - Examples of mosaic phenotypes. (a)** Photograph of an injected G0 male displaying a *white-/ white+* mosaic phenotype on the left eye. **(b)** Photograph of several *ebony* (e1 + e2 + e4) G0 injected animals displaying a range of shading of body pigmentation **(c)** Detail of the bottom two mosquitoes in panel **(b).**

**Figure S2**

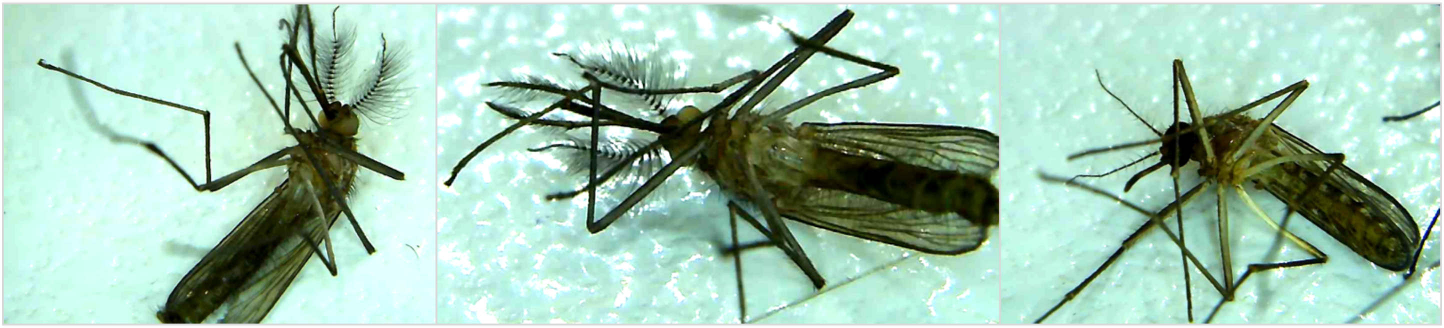

**Figure S2 - *ebony-/white-* phenotype.** Photograph of ebony-white double mutant males (left and middle) compared to a wild-type female (right). Brightness was increased by 20% for all three photographs.

### Figure S3

**a**

```
White

                                   GGATCAAGGGTCTGTCG|GGTGGG
WT-A AGTGTGCGGATACGGTCATCGGAGCGCCCGGACGGATCAAGGGCCTGTCG|GGTGGGGAGCGGAAACGACTGGCGTTCGCGTCGGAAACCCTCACCGATCC
WT-B AGTGTGCGGACACGGTCATCGGAGCGCCAGGACGGATCAAGGGTCTGTCG|GGTGGGGAGAGGAAACGTCTGGCGTTCGCGTCGGAAACACTCACCGATCC
WT-C AGTGTGCCGATACGATCATCGGAGCGCCCGGACGGATTAAGGGACTGTCG|GGTGGGGAGCGGAAACGACTGGCGTTTGCGTCGGAAACCCTCACCGATCC

a(B) AGTGTGCGGACACGGTCATCGGAGCGCCAGGACGGATCAAGGGTCTG---|-GTGGGGAGAGGAAACGTCTGGCGTTCGCGTCGGAAACACTCACCGATCC -4
b(A) AGTGTGCGGATACGGTCATCGGAGCGCCCGGACGGATCAAGGGCCTGTCG|--TGGGGAGCGGAAACGACTGGCGTTCGCGTCGGAAACCCTCACCGATCC -2
c(C) AGTGTGCCGATACGATCATCGGAGCGCCCGGACGGATTAAGGGACTGT--|---GGGGAGCGGAAACGACTGGCGTTTGCGTCGGAAACCCTCACCGATCC -5
d(A) AGTGTGCGGATACGGTCATCGGAGCGCCCGGACGGATCAAGGGCCTGTCG|----GGGAGCGGAAACGACTGGCGTTCGCGTCGGAAACCCTCACCGATCC -4
e(B) AGTGTGCGGACACGGTCATCGGAGCGCCAGGACGGATCAAGGGTCTGT--|---GGGGAGAGGAAACGTCTGGCGTTCGCGTCGGAAACACTCACCGATCC -5
```

**b**

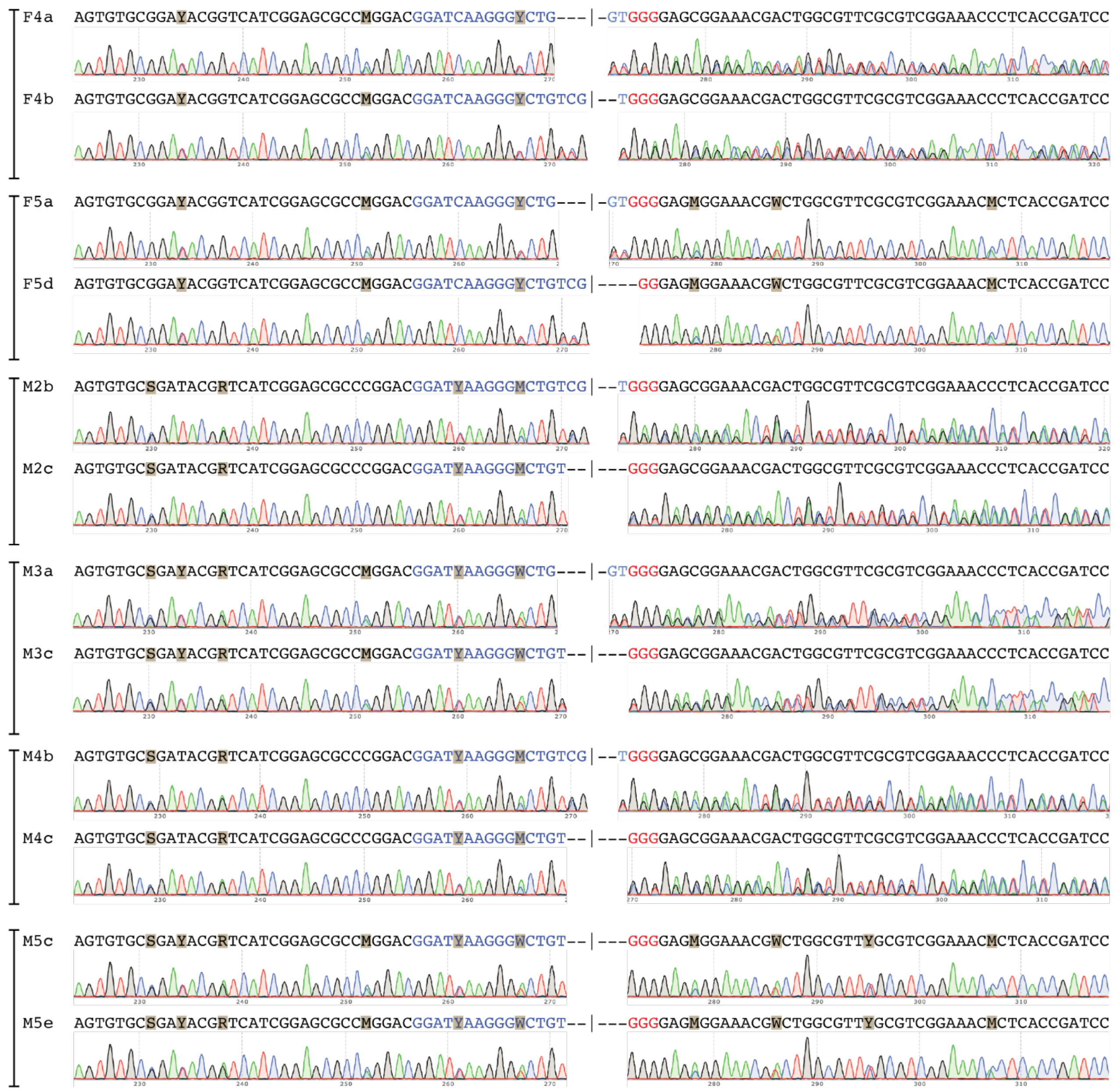


**Figure S3 - Sequencing of *white-* mutant mosquitoes.** (**a**) WT sequences identified, indicated as separate haplotypes labeled as WT-A, WT-B, and WT-C highlighting the SNPs with colored shading matching the traces in (**b**). Below the WT sequences are represented the different indels observed, indicated as a, b, c, d, and e.

Next to each allele, in parentheses, is the letter code for the respective WT allele. The number of deleted bases in each indel is reported on their right. (**b**) Sequences identified for different mosquitoes sequenced, indicated as F (female) or M (male), followed by the number identifier, followed by the allele code (a, b, c, d, e) from panel (**a**). Note: while we have previously identified the sequence for the haplotypes WT-A and WT-B, the sequence of the WT-C allele was obtained by deduction using the sequence of M5 and subtracting the bases from the WT-B allele at the mixed trace positions.

**Figure S4**

**a**

```
kynurenine-hydroxylase (kh)
                                          GGAGTTTAGCGAAACTC|GGTGGG
WT   CAAGCACGGATCGGACGTTGATCAGATCCTGAAGGAGTTTAGCGAAACTC|GGTGGGAGGACGCCCACGCGATCTGCGATTTGGCCATGTACAACTACATC
a    CAAGCACGGATCGGACGTTGATCAGATCCTGAAGGAGTTT----------|---------------ACGCGATCTGCGATTTGGCCATGTACAACTACATC -25
b    CAAGCACGGATCGGACGTTGATCAGATCCTGAAGGAGTTTAGCGAAACTC|-------GGACGCCCACGCGATCTGCGATTTGGCCATGTACAACTACATC -7
c    CAAGCACGGATCGGACGTTGATCAGATCCTGAAGGAGTTTAGCGAAACT-|GGTGGGAGGACGCCCACGCGATCTGCGATTTGGCCATGTACAACTACATC -1
```

**b**

```
F1a  CAAGCACGGATCGGACGTTGATCAGATCCTGAAGGAGTTT----------|---------------ACGCGATCTGCGATTTGGCCATGTACAACTACATC
F1b  CAAGCACGGATCGGACGTTGATCAGATCCTGAAGGAGTTTAGCGAAACTC|-------GGACGCCCACGCGATCTGCGATTTGGCCATGTACAACTACATC

F2a  CAAGCACGGATCGGACGTTGATCAGATCCTGAAGGAGTTT----------|---------------ACGCGATCTGCGATTTGGCCATGTACAACTACATC

F4a  CAAGCACGGATCGGACGTTGATCAGATCCTGAAGGAGTTT----------|---------------ACGCGATCTGCGATTTGGCCATGTACAACTACATC
F4b  CAAGCACGGATCGGACGTTGATCAGATCCTGAAGGAGTTTAGCGAAACTC|-------GGACGCCCACGCGATCTGCGATTTGGCCATGTACAACTACATC

F5a  CAAGCACGGATCGGACGTTGATCAGATCCTGAAGGAGTTT----------|---------------ACGCGATCTGCGATTTGGCCATGTACAACTACATC
F5b  CAAGCACGGATCGGACGTTGATCAGATCCTGAAGGAGTTTAGCGAAACTC|-------GGACGCCCACGCGATCTGCGATTTGGCCATGTACAACTACATC

M1a  CAAGCACGGATCGGACGTTGATCAGATCCTGAAGGAGTTT----------|---------------ACGCGATCTGCGATTTGGCCATGTACAACTACATC
M1c  CAAGCACGGATCGGACGTTGATCAGATCCTGAAGGAGTTTAGCGAAACT-|GGTGGGAGGACGCCCACGCGATCTGCGATTTGGCCATGTACAACTACATC

M2a  CAAGCACGGATCGGACGTTGATCAGATCCTGAAGGAGTTT----------|---------------ACGCGATCTGCGATTTGGCCATGTACAACTACATC
M2b  CAAGCACGGATCGGACGTTGATCAGATCCTGAAGGAGTTTAGCGAAACTC|-------GGACGCCCACGCGATCTGCGATTTGGCCATGTACAACTACATC

M3a  CAAGCACGGATCGGACGTTGATCAGATCCTGAAGGAGTTT----------|---------------ACGCGATCTGCGATTTGGCCATGTACAACTACATC
```

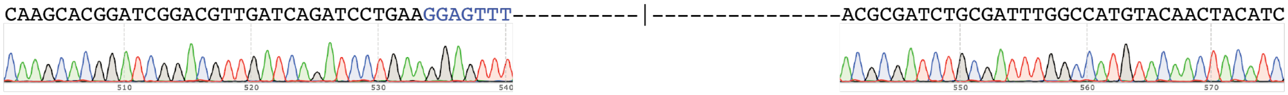

```
M5a  CAAGCACGGATCGGACGTTGATCAGATCCTGAAGGAGTTT----------|---------------ACGCGATCTGCGATTTGGCCATGTACAACTACATC
```

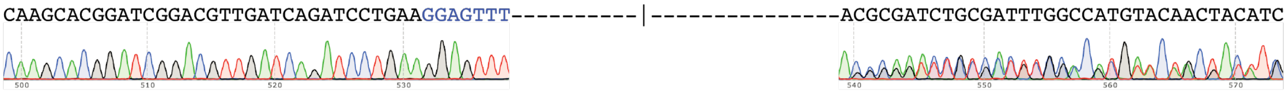

```
M5b  CAAGCACGGATCGGACGTTGATCAGATCCTGAAGGAGTTTAGCGAAACTC|-------GGACGCCCACGCGATCTGCGATTTGGCCATGTACAACTACATC
```

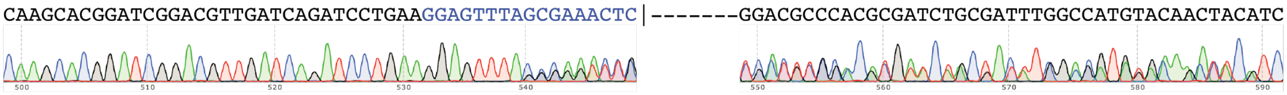

**Figure S4 - Sequencing of *kynurenine-hydroxylase-* mutant mosquitoes.** (**a**) WT sequences identified. Below it, are the different indels observed, indicated as a, b and c, as well as the number of deleted bases on their right. (**b**) Sequences identified for different mosquitoes sequenced, indicated as F (female) or M (male), followed by the number identifier, followed by the allele code (a, b, c) from panel (**a**). For two of the sequenced mosquitoes we report the Sanger sequencing trace, and show how it aligns to either a homozygous or heterozygous mosquito.

### Figure S5

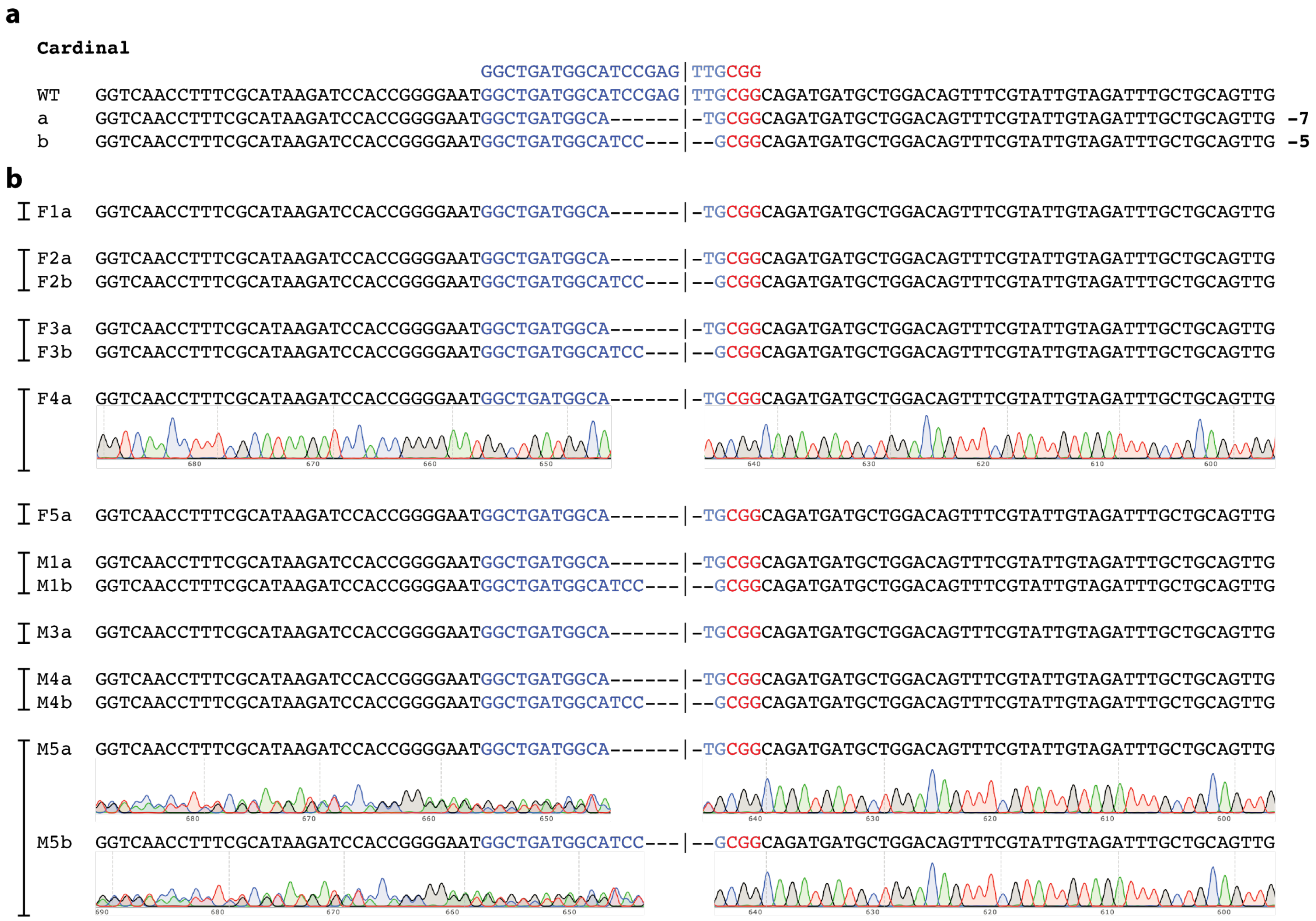


**Figure S5 - Sequencing of *cardinal-* mutant mosquitoes.** (**a**) WT sequences identified. Below it, are the different indels observed, indicated as a and b, as well as the number of deleted bases on their right. (**b**) Sequences identified for different mosquitoes sequenced, indicated as F (female) or M (male), followed by the number identifier, followed by the allele code (a, b) from panel (**a**). For two of the sequenced mosquitoes we report the Sanger sequencing trace, and show how it aligns to either a homozygous or heterozygous mosquito.

### Figure S6

**a**

**Yellow**

```
                                  GGCTGGGCTGAACTTCC|AGTGGG
WT    CTTCCCGGATCCGCTGCGCGGAGACTTCAACGTGGCTGGGCTGAACTTCC|AGTGGGGTGAGGAGGGTATCTTTGGAATGTCGCTGTCGCCTATTCAAGCC
a     CTTCCCGGATCCGCTGCGCGGAGACTTCAACGTGGCTGGGCTGAACTTC-|--TGGGGTGAGGAGGGTATCTTTGGAATGTCGCTGTCGCCGATCCAAGCC -3
```

**b**

```
M1a   CTTCCCGGATCCGCTGCGCGGAGACTTCAACGTGGCTGGGCTGAACTTC-|--TGGGGTGAGGAGGGTATCTTTGGAATGTCGCTGTCGCCGATCCAAGCC
M2a   CTTCCCGGATCCGCTGCGCGGAGACTTCAACGTGGCTGGGCTGAACTTC-|--TGGGGTGAGGAGGGTATCTTTGGAATGTCGCTGTCGCCGATCCAAGCC
M3a   CTTCCCGGATCCGCTGCGCGGAGACTTCAACGTGGCTGGGCTGAACTTC-|--TGGGGTGAGGAGGGTATCTTTGGAATGTCGCTGTCGCCGATCCAAGCC
M4a   CTTCCCGGATCCGCTGCGCGGAGACTTCAACGTGGCTGGGCTGAACTTC-|--TGGGGTGAGGAGGGTATCTTTGGAATGTCGCTGTCGCCGATCCAAGCC
F4a   CTTCCCGGATCCGCTGCGCGGAGACTTCAACGTGGCTGGGCTGAACTTC-|--TGGGGTGAGGAGGGTATCTTTGGAATGTCGCTGTCGCCGATCCAAGCC
```

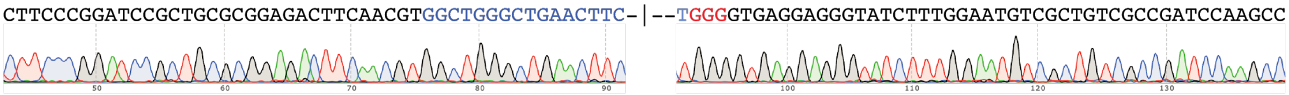


**Figure S6 - Sequencing of *yellow-* mutant mosquitoes.** (**a**) WT sequences identified. Below it, is represented the only indel observed, indicated as a, as well as the number of deleted bases on its right. (**b**) Sequences identified for different mosquitoes sequenced, indicated as F (female) or M (male), followed by the number identifier, followed by the allele code (a) from panel (**a**). For one of the sequenced mosquitoes we report the Sanger sequencing trace, and show how it aligns to the indel observed.

### Figure S7

**a**

```
ebony (gRNA-1)
                                    GGGTTTGACCGGCTGGA|GGTCGG
WT     CAAACCGGACACGAACAGTTCACCGATCTCACCGGGTTTGACCGGCTGGA|GGTCGGGAGTTAGGATGTATATCACAGTGTTGTCCAGGGGGTACCCGATA
a      CAAACCGGACACGAACAGTTCACCGATCTCACCGGGTTTGACC-------|GGTCGGGAGTTAGGATGTATATCACAGTGTTGTCCAGGGGGTACCCGATA -7
b      CAAACCGGACACGAACAGTTCACCGATCTCACCGGGTTTGACCGGCT---|----GGGAGTTAGGATGTATATCACAGTGTTGTCCAGGGGGTACCCGATA -7
```

**b**

```
M1a    CAAACCGGACACGAACAGTTCACCGATCTCACCGGGTTTGACC-------|GGTCGGGAGTTAGGATGTATATCACAGTGTTGTCCAGGGGGTACCCGATA
M2b    CAAACCGGACACGAACAGTTCACCGATCTCACCGGGTTTGACCGGCT---|----GGGAGTTAGGATGTATATCACAGTGTTGTCCAGGGGGTACCCGATA
```

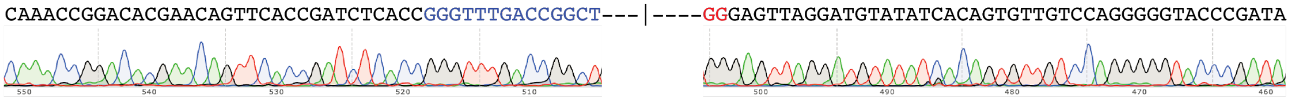

```
M3a    CAAACCGGACACGAACAGTTCACCGATCTCACCGGGTTTGACC-------|GGTCGGGAGTTAGGATGTATATCACAGTGTTGTCCAGGGGGTACCCGATA
```

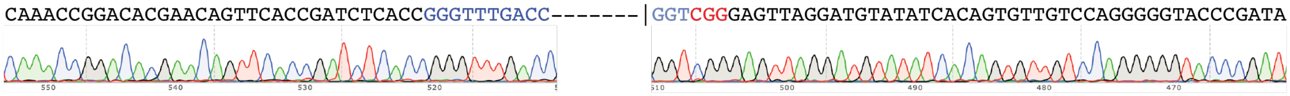

```
M4a    CAAACCGGACACGAACAGTTCACCGATCTCACCGGGTTTGACC-------|GGTCGGGAGTTAGGATGTATATCACAGTGTTGTCCAGGGGGTACCCGATA
```

**c**

```
ebony (gRNA-2)
                                    GGTCACGGAGGCAAAGT|CTCCGG
WT     ATCCGTACGACCTTCGTAGTACACGCAGCCCTTGGTCACGGAGGCAAAGT|CTCCGGTGCGGTACATCCTTGAGTAAATGAGGTCAACAGCCAGTGGATTG
a      ATCCGTACGACCTTCGTAGTACACGCAGCCCTTGGTCACGGAGGCAAAGT|--CCGGTGCGGTACATCCTTGAGTAAATAAGGTCAACAGCCAGTGGATTG -2
b      ATCCGTACGACCTTCGTAGTACACGCAGCCCTTGGTCACGGAGGCAAAGT|-------GCGGTACATCCTTGAGTAAATAAGGTCAACAGCCAGTGGATTG -7
```

**d**

```
M1a    ATCCGTACGACCTTCGTAGTACACGCAGCCCTTGGTCACGGAGGCAAAGT|--CCGGTGCGGTACATCCTTGAGTAAATAAGGTCAACAGCCAGTGGATTG
M2b    ATCCGTACGACCTTCGTAGTACACGCAGCCCTTGGTCACGGAGGCAAAGT|-------GCGGTACATCCTTGAGTAAATAAGGTCAACAGCCAGTGGATTG
```

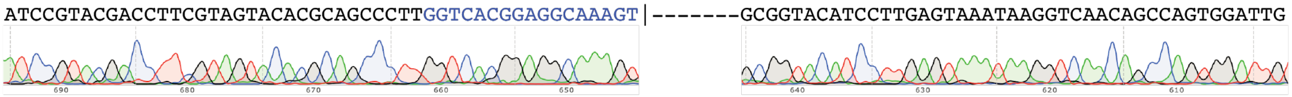

```
M3a    ATCCGTACGACCTTCGTAGTACACGCAGCCCTTGGTCACGGAGGCAAAGT|--CCGGTGCGGTACATCCTTGAGTAAATAAGGTCAACAGCCAGTGGATTG
```

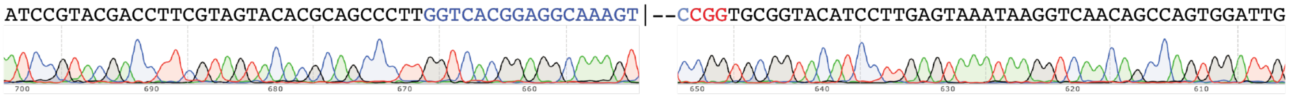

```
M4a    ATCCGTACGACCTTCGTAGTACACGCAGCCCTTGGTCACGGAGGCAAAGT|--CCGGTGCGGTACATCCTTGAGTAAATAAGGTCAACAGCCAGTGGATTG
```

**Figure S7 - Sequencing of *ebony-* mutant mosquitoes.** Sequences of four male mosquitos, at the *e1*-gRNA (**a**, **b**) and *e2*-gRNA (**c**, **d**) locations.(**a, c**) WT sequences identified at the *e1*-gRNA and *e2*-gRNA. Below them are represented the indels observed, indicated as “a” or “b”, as well as the number of deleted bases on their right. (**b, d**) Sequences identified for different mosquitoes sequenced, indicated as F (female) or M (male), followed by the number identifier, followed by the allele code (a or b) from panel (**a** or **c**). For two of the sequenced mosquitoes we report the Sanger sequencing trace, and show how it aligns to the indels observed.

### Figure S8

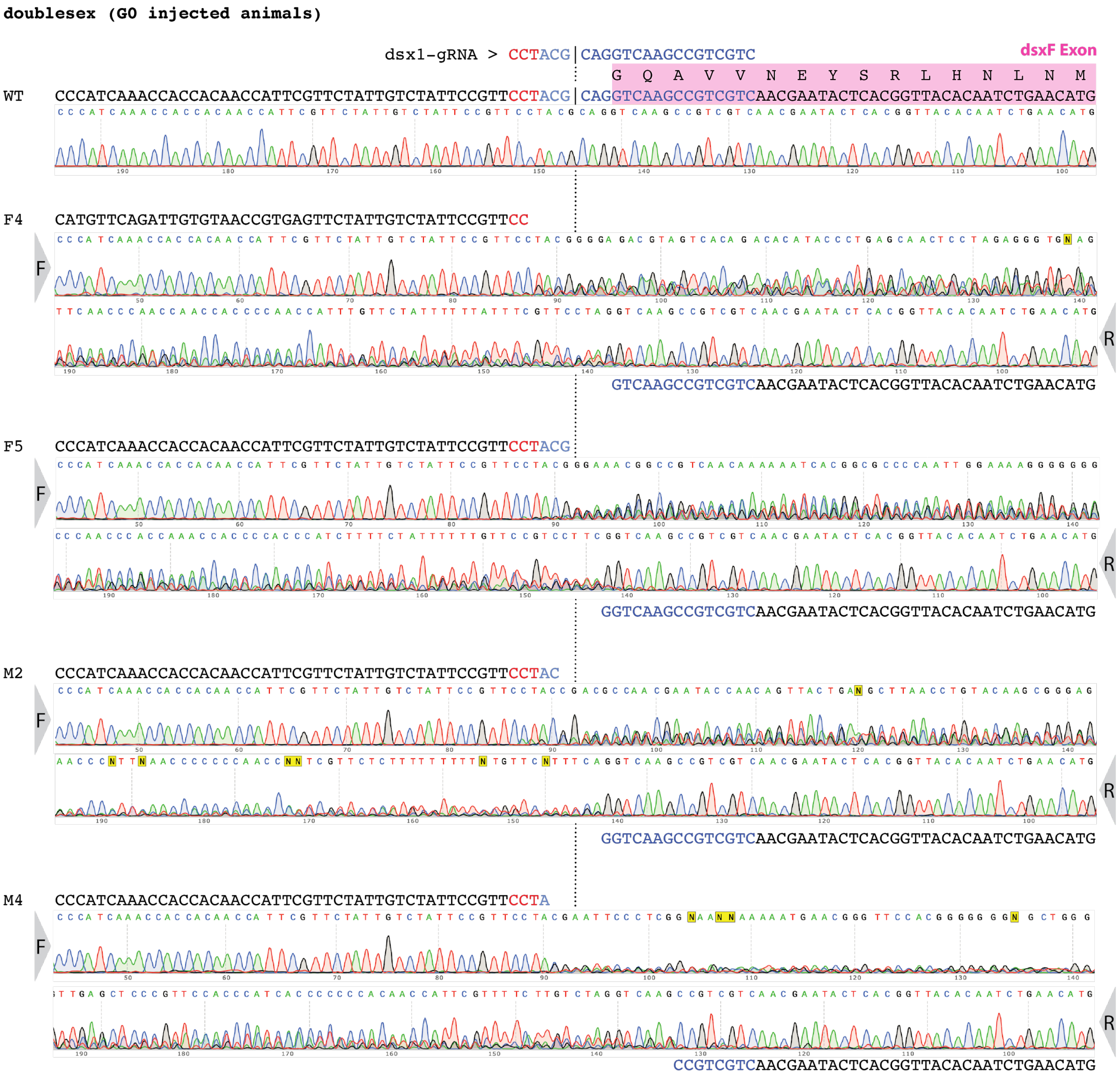


**Figure S8 - Sequencing of *doublesex* G0 injected mosquitoes.** In order, sequencing traces of wild-type and two female and two male G0 injected mosquitos displaying mixed traces beginning around the cut site. For each mosquito a forward and a reverse read are displayed, indicated with grey arrows on the side. In pink shading is represented the dsxF exon position.

**Figure S9**

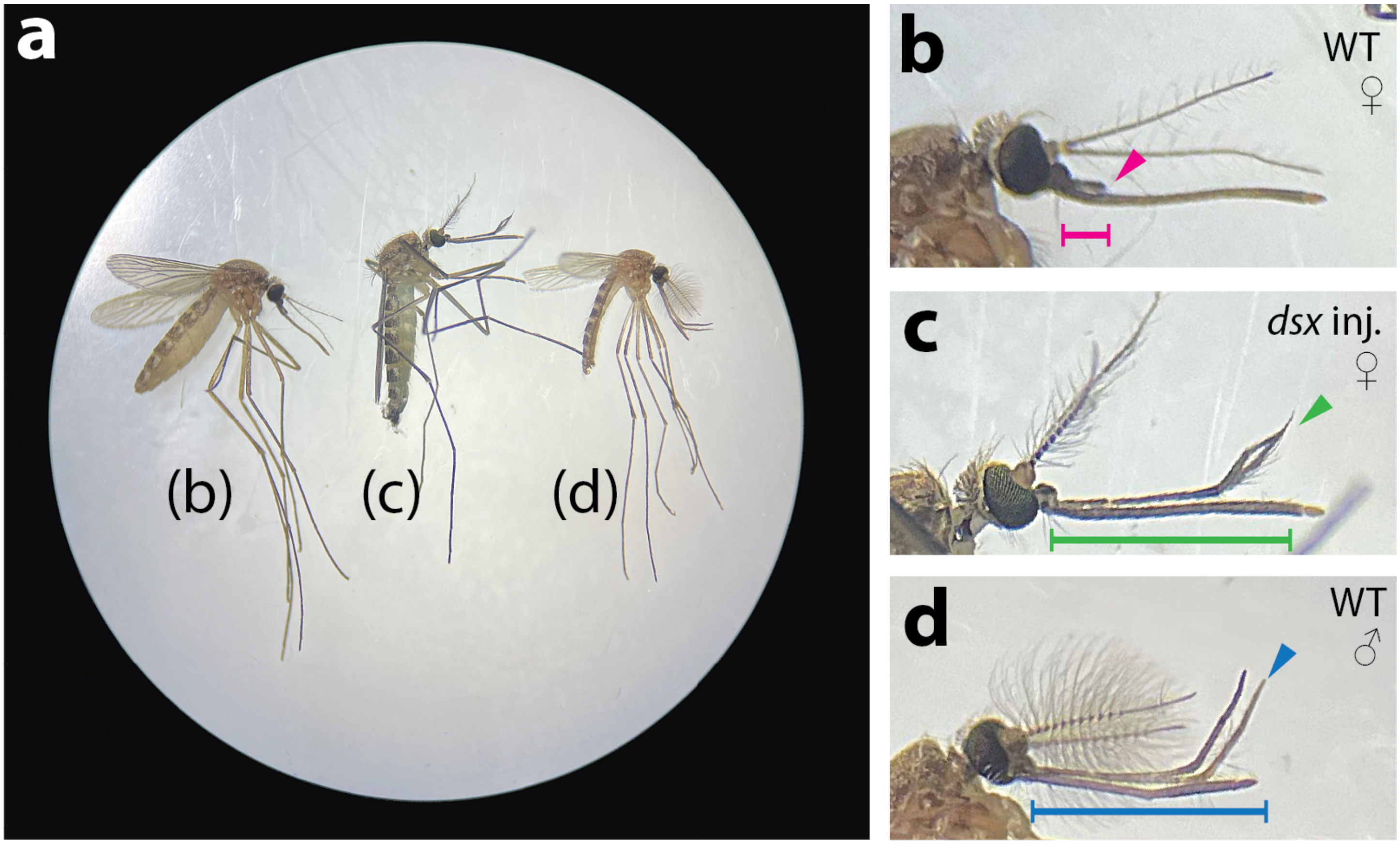


**Figure S9 - Phenotype of *doublesex* editing in the Hawai'i sourced line. (a)** Microscope image of **(b)** a wild-type female, **(c)** a female from the injection of Cas9/gRNA targeting the dsxF transcript, and **(d)** a wild-type male. **(b-d)** Magnifications of the mosquitoes in **(a)**. **(b)** Wild-type female with short MPs (magenta bar and arrow); **(c)** *dsx*-injected G0 female showing elongated MPs (green bar and arrow). **(d)** Wild-type male with long maxillary palps (MP) curved upwards (blue bar and arrow).
